# Potential of Ilhéus virus to emerge

**DOI:** 10.1101/2023.09.08.556909

**Authors:** Kenneth S. Plante, Jessica A. Plante, Sasha R. Azar, Divya P. Shinde, Dionna Scharton, Alice F. Versiani, Natalia Ingrid Oliveira da Silva, Taylor Strange, Lívia Sacchetto, Eric B. Fokam, Shannan L. Rossi, Scott C. Weaver, Rafael E. Marques, Mauricio L. Nogueira, Nikolaos Vasilakis

## Abstract

Ilhéus virus (ILHV)(*Flaviviridae: Orthoflavivirus*) is an arthropod-borne virus (arbovirus) endemic to Central and South America and the Caribbean. First isolated in 1944, most of our knowledge derives from surveillance and seroprevalence studies. These efforts have detected ILHV in a broad range of mosquito and vertebrate species, including humans, but laboratory investigations of pathogenesis and vector competence have been lacking. Here, we develop several immune intact murine models that closely recapitulate human neuroinvasive disease with strain- and age-specific virulence, as well as a uniformly lethal immunocompromised model. Replication kinetics in several vertebrate and invertebrate cell lines demonstrate that ILHV is capable of replicating to high titers in a wide variety of potential host and vector species. Lastly, vector competence studies provide strong evidence for efficient infection of and potential transmission by *Aedes* species mosquitoes, despite ILHV’s phylogenetically clustering with *Culex* vectored flaviviruses, suggesting ILHV is poised for emergence in the neotropics.

**Teaser:** Murine models of ILHV mimic human disease, and *Aedes* species of mosquitoes are highly susceptible to infection and dissemination.

## Introduction

Ilhéus virus (ILHV) is a neuroinvasive neglected tropical pathogen of the Ntaya antigenic group in the family *Flaviviridae*, genus *Orthoflavivirus*, and has the potential to pose a threat to human health. The virus was originally isolated in 1944 from *Psorophora* and *Aedes spp*. mosquitoes in Ilhéus, State of Bahia, Brazil (*1, 2*). In the following years, the virus was detected in several mosquito spp., including *Culex, Coquillettidia, Haemagogus, Ochlerotatus, Sabethes*, and *Trichoprosopon*. ILHV is believed to be maintained in an enzootic cycle between arboreal mosquitoes, primarily *Culex*. spp., and birds (*3-9*). The virus has been commonly detected by viral isolation or PCR-based methods in various bird species and humans throughout Central and South America and the Caribbean (*10-17*). However, serologic surveys indicate that ILHV may infect a wide range of animals including non-human primates, horses, coatis, rodents, bats, tortoises, and sloths (*7, 16, 18-32*). This broad circulation in the neotropics, accompanied by human expansion into previously unpopulated or sparsely populated regions, makes ILHV a threat for spillover events into human populations.

To date, ILHV has not been associated with any large epidemics, although a close relative, Rocio virus (ROCV), was responsible for an outbreak in 1976 that infected well over 1,000 people with a 13% case fatality rate (*33*). ILHV is the etiologic agent of Ilhéus fever, and has been sporadically reported in Bolivia, Brazil, Columbia, Ecuador, French Guiana, Panama, and Trinidad (*11-15, 32, 34*). Ilhéus fever generally presents as an undifferentiated febrile illness that can progress to neurological involvement (*13, 14, 32, 34-36*), and reviewed in (*37*). In 2017, a retrospective study of patients with central nervous system impairment in São Paulo, Brazil detected ILHV genomic material in the cerebrospinal fluid of an elderly male patient with right hemiplegia, dysarthria, aphasia, and intraparenchymal hemorrhage (*38*). This is believed to be the first report of fatal ILHV infection in a human.

Herein, we leveraged the extensive collection of the World Reference Center for Emerging Viruses and Arboviruses (WRCEVA) to better understand ILHV’s potential to emerge and cause epidemics. Multiple strains of ILHV were characterized to represent the maximum degree of geographic, temporal, and genetic diversity within the viral species (*39*). Replication kinetics assessed in immortalized cell lines from multiple mammalian and mosquito spp. supports the serological evidence for ILHV circulation in a wide array of host and vector species. We established an immune-intact murine model which mimics the neuroinvasive phenotype of ILHV in humans in an age-dependent manner. The immunocompromised murine model that we established develops high viremias and is uniformly lethal, rendering it useful for stringent testing of potential therapeutic or vaccine interventions as well as for vector competence studies. Our vector competence studies indicate that, although ILHV is a neurotropic flavivirus that utilizes birds as primary amplification and reservoir hosts and groups phylogenetically with other *Culex*-vectored flaviviruses (*40, 41*), the *Aedes* spp. mosquitoes were far more susceptible to infection in a laboratory setting. These studies increase our understanding of this neglected tropical pathogen and create a pathway for further studies of this potential emergent and underreported threat to human health.

## Results

### Viral Replication in Cultured Cells

The replication kinetics of eight ILHV strains (**Table S1**) were characterized in seven cell lines. Four of the cell lines are mammalian: Vero (*Cercopithecus aethiops* kidney epithelial), Huh-7 (*Homo sapiens* hepatocyte), BHK (*Mesocricetus auratus* kidney fibroblast), and OK (*Didelphis marsupialis virginiana* kidney epithelial). Three of the cell lines are derived from the larva of *Ae. albopictus* mosquitoes: C6/36, C7/10, and U4.4. All eight ILHV strains exhibited robust replication in all seven cell lines (**Fig. 1**). In mammalian cells (**Fig. 1A-D**), ILHV rapidly reached its peak titer of 7.14-7.83 log_10_ FFU (focus forming units)/ml, most often at three days post-infection (DPI) (range: 2-4 DPI). The post-peak decline in ILHV titers corresponded with the onset of cytopathic effect (CPE) in the cells (**Figs. S1-S4**). In contrast, ILHV replication in the three *Ae. albopictus* cell lines was characterized by delayed peak titers without an appreciable post-peak decline (**Fig. 1E-G**). This is consistent with visual observations of the infected mosquito cell monolayers, which exhibited clumping and syncytia formation but not the destructive CPE seen in the mammalian cell lines (**Figs. S5-S7**). The two RNAi-deficient *Ae. albopictus* cell lines, C6/36 and C7/10, supported peak ILHV titers of 6.81-8.06 log_10_ FFU/ml and 8.20-9.54 log_10_ FFU/ml, respectively. In contrast, the RNAi-competent U4.4 line only supported peak ILHV titers of 5.73-6.45 log_10_ FFU/ml. Strain based variation in ILHV replication was generally non-significant or weakly significant, especially at or prior to peak titer. Significant differences were more common at later timepoints, when degraded or destroyed cell monolayers are more likely to have a prominent impact (**Figs. S1-S7 and Table S2**). These results support the serologic and mosquito survey evidence for the ability of ILHV to productively infect a broad range of potential host and vector species.

**Fig. 1.**
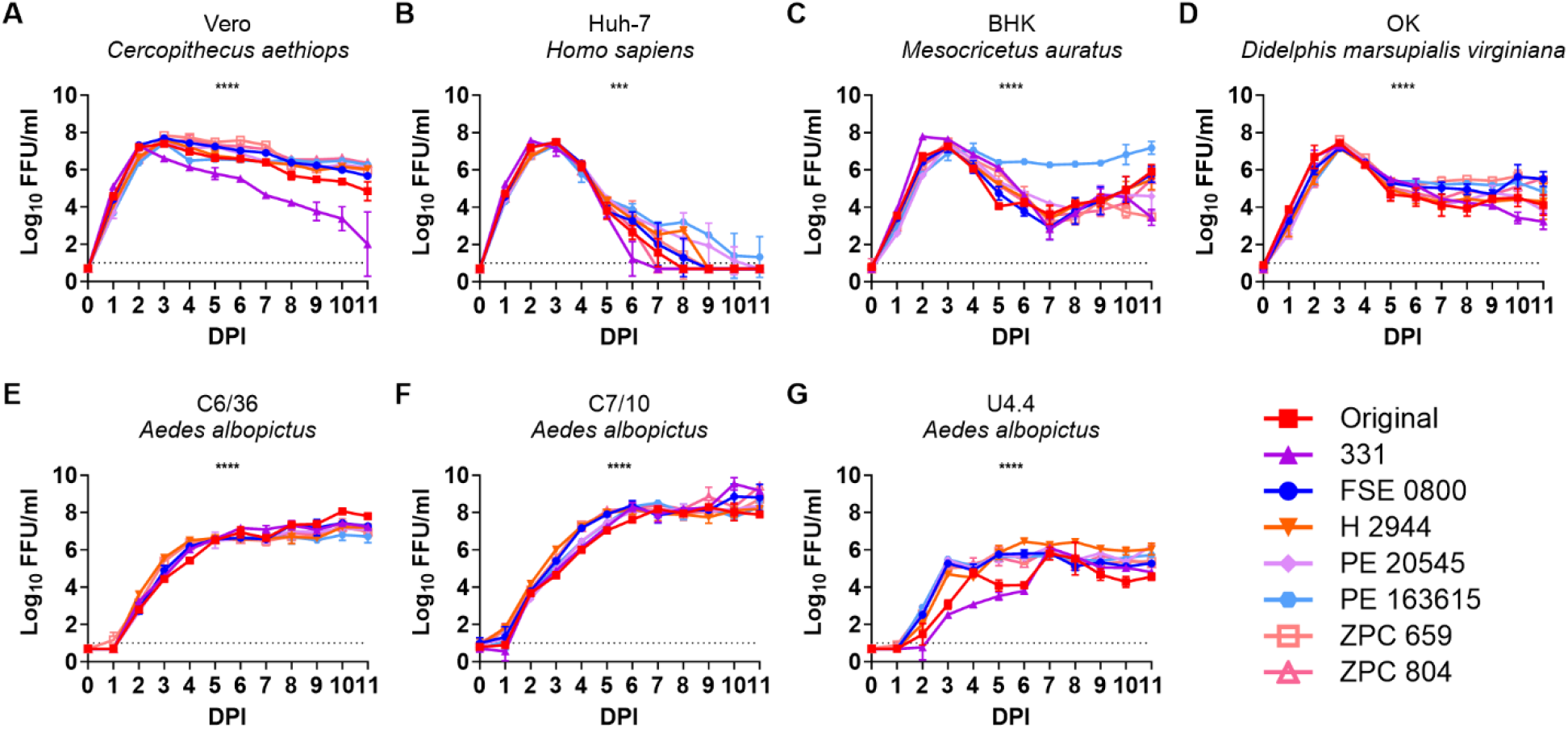
ILHV replicates to high titers in cultured cells from a variety of mammalian and mosquito species. (**A**) Vero, (**B**) Huh-7, (**C**) BHK, (**D**) OK, (**E**) C6/36, (**F**) C7/10, and (**G**) U4.4 cells were infected with eight strains of ILHV at a multiplicity of infection of 0.01. Samples were collected daily for 11 DPI, and viral titer was determined by focus forming assay in Vero cells. Data is representative of three independent replicates from a single experiment. Colors represent the ILHV strain, symbols represent the mean, and error bars represent the standard deviation. Dotted lines represent the lower limit of detection. Samples with no detectable ILHV are represented as half the lower limit of detection. Log_10_-transformed titers were analyzed by two-way ANOVA; the significance of overall ILHV strain-based variation is presented within each panel, and the daily strain by strain comparisons with Tukey’s multiple comparisons tests are presented in Table S2. ns = p≥0.05, * = p<0.05, ** = p<0.01, *** = p<0.001, **** = p<0.0001.

### Establishment of Murine Models of Pathogenesis

There is no well-characterized small animal model for ILHV infection and disease. Therefore, four-week-old male CD-1 mice were infected with 5.0 log_10_ FFU of either ILHV FSE 0800 or ILHV Original, or mock-infected with PBS, via the intraperitoneal (IP) route (**Fig. 2A-B**). The FSE 0800 and Original strains of ILHV represent the temporal extremes of our available ILHV panel (**Table S1**); FSE 0800 was isolated in 2004 and has been passaged four times in Vero cells and is considered as close to non-adapted, whereas Original was isolated in 1944 and has been passaged 29 times in suckling mice and twice in Vero cells, and thus is considered mouse-adapted. ILHV Original was universally lethal, with an average survival time of 5.6 days and all mice succumbing to infection by eight DPI. ILHV FSE 0800, in contrast, was only lethal in 10% of mice. Mice that succumbed to ILHV infection exhibited hunched posture (8/11), ruffled fur (6/11), lethargy (6/11), tremors (5/11), and grimace (3/11), with onset either the day of death or one day prior. ILHV Original-infected mice lost a significant amount of weight compared to mock-infected mice on days four through eight post-infection, corresponding to the timeframe during which ILHV Original-infected mice died. The ILHV Original-infected mice also lost significantly more weight than the ILHV FSE 0800-infected mice during this time except for day seven, the day on which a single ILHV FSE 0800 mouse succumbed to infection.

**Fig. 2.**
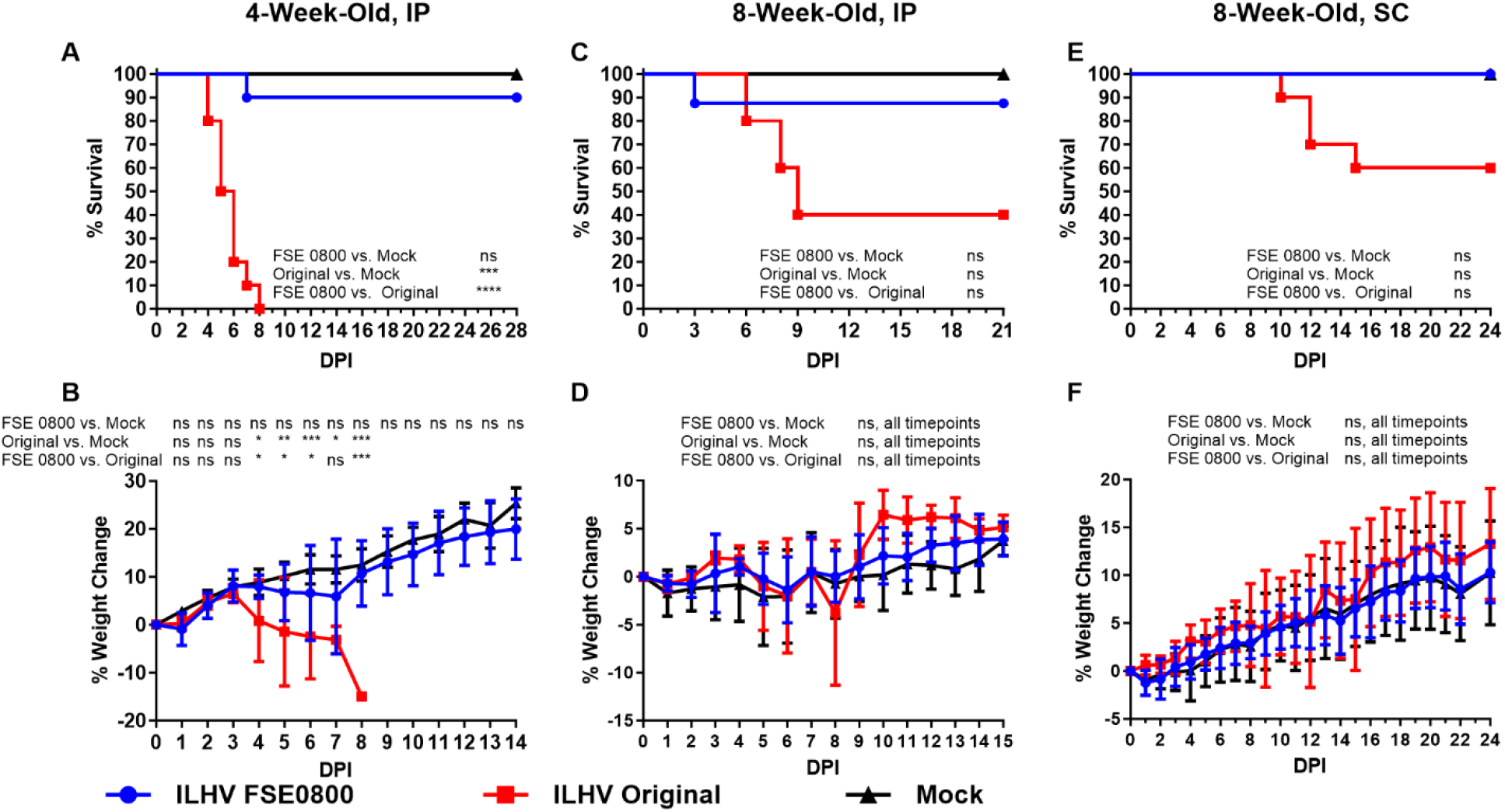
Mortality and weight loss in a CD-1 model is highly dependent on ILHV strain and mouse age. (**A-B**) four-or (**C-F**) eight-week-old male CD-1 mice were infected via either the (**A-D**) intraperitoneal or (**E-F**) subcutaneous route with 5.0 log_10_ FFU of ILHV FSE 0800, ILHV Original, or a PBS-only mock control. (**A, C, E**) Survival rates and (**B, D, F**) weight change are reported. Colors represent the ILHV strain or mock cohort. For weight loss, symbols represent the mean and error bars represent the standard deviation. The (**A**-**B**) four-week-old IP route and (**E-F**) eight-week-old SC route was assessed in n=10 ILHV FSE 0800- and ILHV Original-infected mice and in n=5 mock-infected mice. The (**C**-**D**) eight-week-old IP route was assessed in n=5 ILHV FSE 0800- and ILHV Original-infected mice and in n=2 mock-infected mice. Data are representative of a single experiment. Survival was assessed by the log-rank (Mantel-Cox) test, with the Holm-Sidak correction for multiple comparisons. Weight changes were assessed by two-way ANOVA with Tukey’s correction for multiple comparisons, except for days 9-14 in the four-week-old model (**B**) which required multiple t-tests with the Holm-Sidak correction due to the presence of two rather than three cohorts. ns = p≥0.05, * = p<0.05, ** = p<0.01, *** = p<0.001, **** = p<0.0001.

Eight-week-old male CD-1 mice were also infected with 5.0 log_10_ FFU of either ILHV FSE 0800 or ILHV Original, or mock-infected with PBS, via either the IP route (**Fig. 2C-D**) or the subcutaneous (SC) route (**Fig. 2E-F**) to assess the potential of the CD-1 mouse model in future studies requiring an older mouse, such as the challenge phase of a vaccine study. ILHV Original was again more virulent than ILHV FSE 0800 in both models, although this difference did not meet the threshold of significance with the cohort sizes utilized. Whereas ILHV Original was 100% lethal in four-week-old IP-infected mice, it was only 60% lethal in eight-week-old IP-infected mice and 40% lethal in eight-week-old SC-infected mice. Time to death was longer in the older models, with ILHV Original resulting in an average survival time of 7.7 days following IP injection and 12.3 days following SC injection. Neither strain of ILHV resulted in significant weight loss in the eight-week-old model. Visible signs of illness were similar to those observed in the four-week-old CD-1 mouse model. In addition to the eight-week-old CD-1 mouse model, six-to eight-week-old A129 mice, of both sexes, were injected SC with 5.0 log_10_ FFU of either ILHV FSE 0800 or ILHV Original, or mock-infected with PBS (**Fig. S8**). This *IFNAR*^-/-^ model was rapidly and uniformly lethal following ILHV infection with either strain, with 100% of mice infected with either ILHV strain succumbing on day three post-infection.

Tropism was assessed in the four-week-old CD-1 model at two and four DPI (**Fig. 3**). Viremia was modest in magnitude and duration (**Fig. 3A**). At one DPI, ILHV FSE 0800 and ILHV Original reached mean titers of 3.5 log_10_ FFU/ml and 3.0 log_10_ FFU/ml, respectively, with all subjects above the limit of detection. Viremia for both strains was below the limit of detection at days two through four post-infection with the exception of a single ILHV Original mouse from the day two and the day three cohorts being viremic at the limit of detection on those days. ILHV Original was highly neurotropic, reaching 3.2 log_10_ FFU/g in the brain at two DPI and 9.1 log_10_ FFU/g at four DPI (**Fig. 3B**). ILHV FSE 0800, on the other hand, was not detected in the brain on day two and reached only 4.5 log_10_ FFU/g on day four. Similar results were observed in the spinal cord and the eye (**Fig. 3C-D**). Virus was detected at low levels (2.4-2.9 log_10_ FFU/g) in the spleen of 60% of ILHV FSE 0800-infected mice at days two and four post-infection, and at 60% and 20% of ILHV Original-infected mice at days two and four post-infection, respectively (**Fig. 3E**). Viral load in the kidneys was similarly low in magnitude and prevalence (**Fig. 3F**). No virus was detected in the liver (**Fig. 3G**). The virus was present at low levels in lung and heart tissue following both ILHV FSE 0800 and ILHV Original infection, with the magnitude and prevalence decreasing from day two to day four in both tissues for each strain (**Fig. 3H-I**). Similar titers were detected in somatic muscle tissue, although there was no trend toward decreasing titer from day two to day four (**Fig. 3J**). Infectious virus was also detected in the testes (**Fig. 3K**). ILHV Original generated relatively low titers (2.1-2.4 log_10_ FFU/g) in 80% of mice on day two and in 40% of mice on day four. ILHV FSE 0800 was detected in 20% of the mice at day two and 80% of the mice at day four. The majority of ILHV FSE 0800-positive mice had similar viral loads (2.3-3.0 log_10_ FFU/g) compared to the ILHV Original-positive mice, but one ILHV FSE 0800 mouse from day four had 7.6 log_10_ FFU/g in its testes. A parallel tropism study was conducted on older (ten-week-old) CD-1 mice (**Fig. S9**). Viremia once again peaked at one DPI and essentially disappeared by two DPI. However, viral loads in the solid organs were generally decreased both in magnitude and in the percentage of virus-positive subjects compared to the four-week-old mice.

**Fig. 3.**
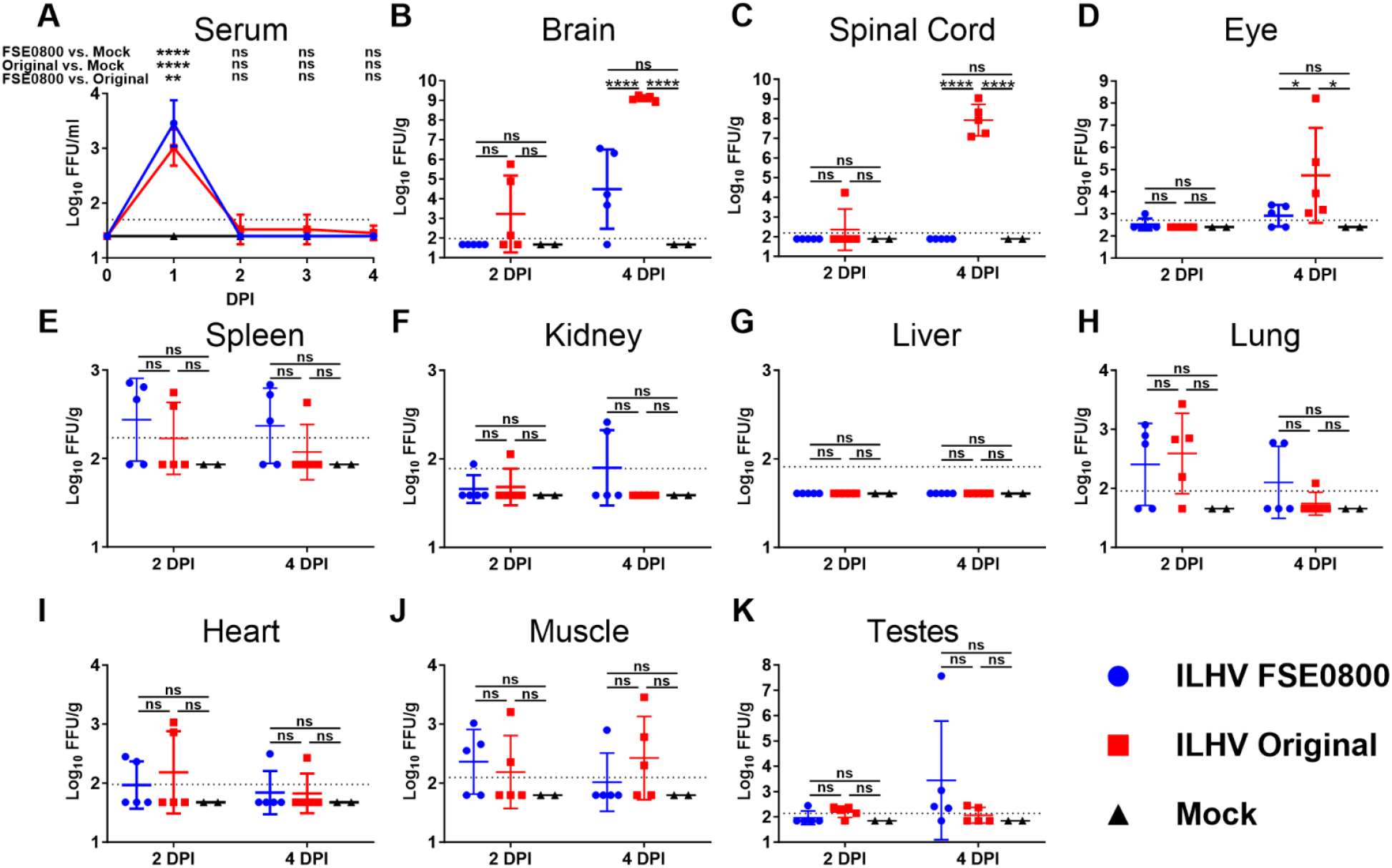
ILHV replicates in multiple murine tissues. Four-week-old male CD-1 mice were infected via the intraperitoneal route with 5.0 log_10_ FFU of ILHV FSE 0800 (n=10), ILHV Original (n=10), or a PBS-only mock control (n=4). Blood was collected from all extant subjects via the retro-orbital route at 1 DPI and 3 DPI. Half of the subjects from each cohort were euthanized at 2 DPI, and the remaining half were euthanized at 4 DPI. At the time of euthanasia, blood was collected prior to PBS perfusion, after which solid organs were collected. Infectious ILHV was detected by focus forming assay in Vero cells. Data are representative of a single experiment. For (**A**) serum samples, symbols represent the mean and error bars represent the standard deviation. For (**B-K**) solid tissues, symbols represent individual subjects, midline bars represent the mean, and error bars represent the standard deviation. Dotted lines represent the lower limit of detection. Samples with no detectable ILHV are represented as half the lower limit of detection. Log_10_-transformed titers were analyzed by two-way ANOVA with Tukey’s multiple comparison’s test. ns = p≥0.05, * = p<0.05, ** = p<0.01, *** = p<0.001, **** = p<0.0001.

### Vector Competence

To establish the vector competence of ILHV, *Culex* spp. and *Aedes* spp. mosquitoes were allowed to feed on viremic A129 mice infected with either ILHV FSE 0800 or ILHV Original. Engorged mosquitoes were maintained for an extrinsic incubation period (EIP) of seven or 14 days, at which times bodies, legs, and saliva were harvested and the presence or absence of ILHV was assessed by cytopathic effect assay in Vero cells with immunostaining. The Sebring colony of *Cx. quinquefasciatus* was highly refractory to both strains of ILHV (**Table 1**). The Salvador colony of *Cx. quinquefasciatus* was also generally poorly infected by ILHV; while the FSE 0800 strain infected 35.7% of the bodies, only a single mosquito had a disseminated and transmissible infection. The *Cx. tarsalis* mosquitoes were more susceptible than the *Cx. quinquefasciatus* mosquitoes. ILHV FSE 0800 was more infectious than ILHV Original in *Cx. tarsalis*, but once again the virus was poorly disseminated; of the 94.1% of *Cx. tarsalis* with ILHV FSE 0800-infected bodies, only 68.8% developed a disseminated infection as assessed by virus in the legs and no mosquitoes became able to transmit the virus as assessed by the absence of virus in the saliva.

**Table 1.**
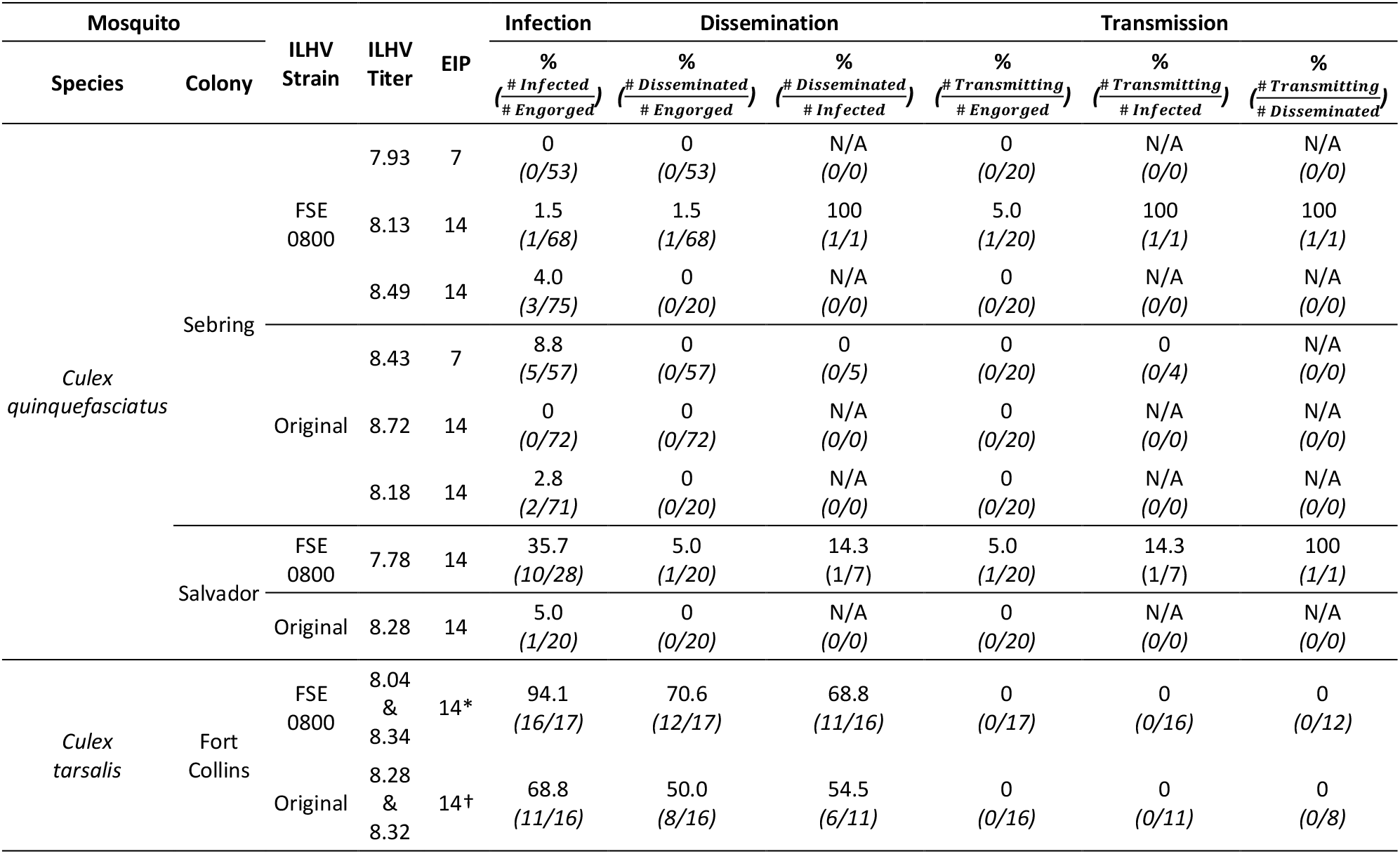
ILHV is poorly disseminated in *Culex* spp. mosquitoes. *Culex* spp. mosquitoes were permitted to feed upon A129 mice previously infected with either ILHV FSE 0800 or ILHV Original. Engorged mosquitoes were maintained for an EIP of seven or 14 days. Bodies, legs, and saliva were collected and the presence of ILHV was detected by infectious assay with immunostaining in Vero cells. ILHV titers detected in the sera of A129 mice immediately following mosquito feeding are reported in FFU/ml. Infection is defined as ILHV in the body, dissemination is defined as ILHV in the legs, and transmission is defined as ILHV in the saliva. Data are representative of a single experiment. ^*^ One mosquito in the cohort had a negative body and positive legs † Two mosquitoes in the cohort had negative bodies and positive legs

In contrast, *Aedes* spp. mosquitoes were highly susceptible to infection by ILHV (**Table 2**). ILHV FSE 0800 was generally more infectious than ILHV Original in the *Aedes* spp. mosquitoes, like the pattern seen in the more refractory *Culex* mosquitoes. In *Ae. aegypti*, ILHV FSE 0800 infected over 90% of bodies following an EIP of seven or 14 days in both the Iquitos and Salvador colonies. ILHV Original infected 79-81% of bodies after seven days and 91-96% of bodies after 14 days. ILHV FSE 0800 disseminated to the legs of infected *Ae. aegypti* quickly and at a high rate, with 98-100% of infected mosquitoes developing a disseminated infection by day seven. ILHV Original disseminated in only 33-43% of infected mosquitoes by day seven but reached 75-100% dissemination by day 14. In further contrast to the infection of *Cx*. mosquitoes, *Ae. aegypti* had virus present in the saliva of 40-100% of FSE 0800-infected mosquitoes and 25-75% of Original-infected mosquitoes with disseminated infections. The results from *Ae. albopictus* were broadly like those of *Ae. aegypti*. ILHV FSE 0800 infected and disseminated to 95-100% of mosquitoes and was present in the saliva of 80-100% of *Ae. albopictus* mosquitoes by day 14. ILHV Original similarly infected 95-100% of bodies but lagged somewhat in dissemination to the legs, infecting only 44-55% of legs on day seven but reaching 70-100% of legs on day 14. Interestingly, although both *Ae. aegypti* and *Ae. albopictus* had high rates of ILHV in the bodies and legs by day seven, only *Ae. aegypti* had ILVH in the saliva by day seven, indicating a more rapid onset of transmission competence in *Ae. aegypti* than in *Ae. albopictus*. This indication that ILHV likely utilizes *Aedes* spp. rather than *Culex* spp. as its primary vector was broadly reflected in the titers of ILHV detected in the bodies, legs, and saliva of *Ae. aegypti, Ae. albopictus*, and *Cx. tarsalis* mosquitoes (**Fig. S10**).

**Table 2.**
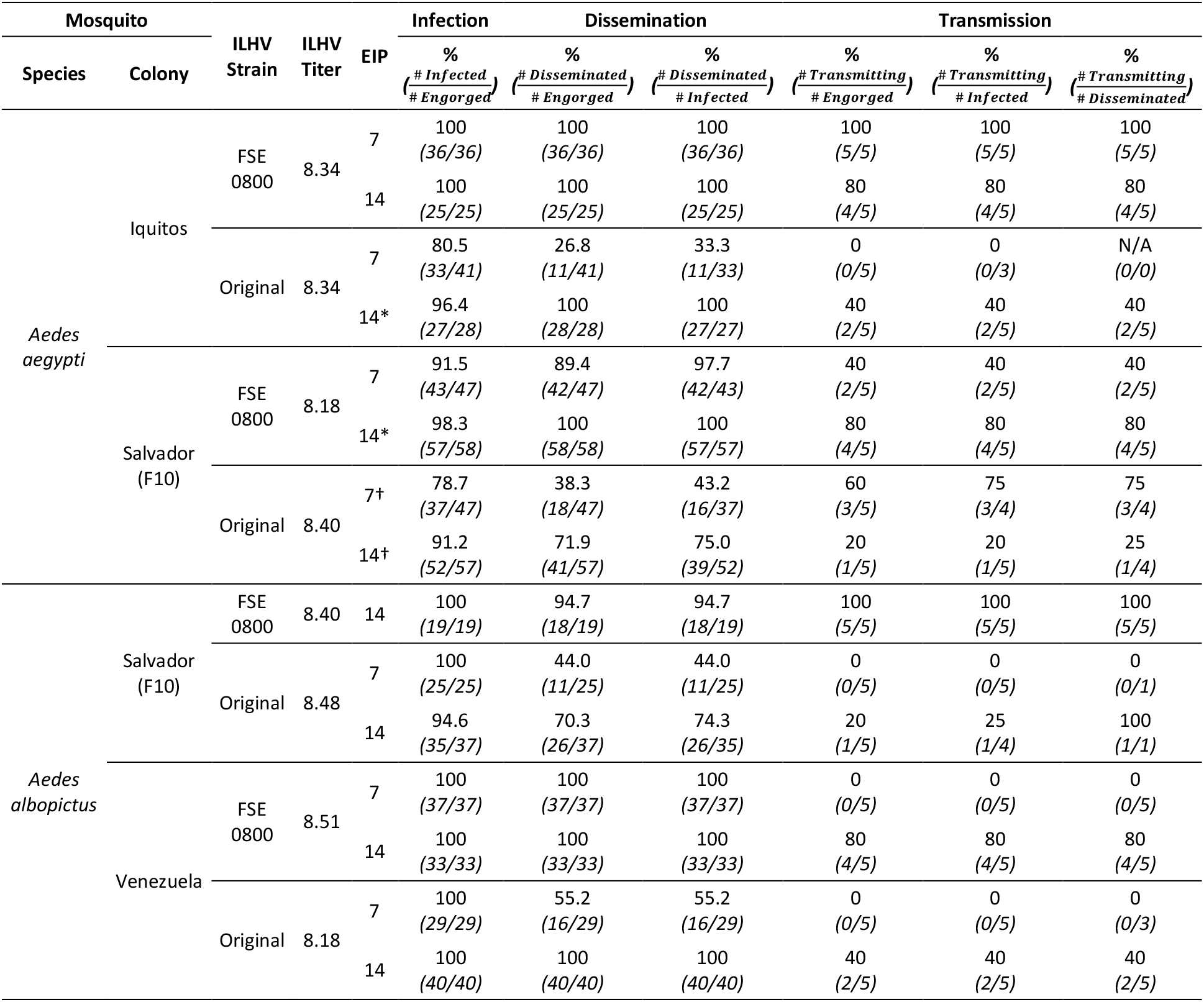
ILHV is efficiently disseminated in *Aedes* spp. mosquitoes. *Aedes* spp. mosquitoes were permitted to feed upon A129 mice previously infected with either ILHV FSE 0800 or ILHV Original. Engorged mosquitoes were maintained for an EIP of seven or 14 days. Bodies, legs, and saliva were collected and the presence of ILHV was detected by infectious assay with immunostaining in Vero cells. ILHV titers detected in the sera of A129 mice immediately following mosquito feeding are reported in FFU/ml. Infection is defined as ILHV in the body, dissemination is defined as ILHV in the legs, and transmission is defined as ILHV in the saliva. Data are representative of a single experiment. ^*^ One mosquito in the cohort had a negative body and positive legs † Two mosquitoes in the cohort had negative bodies and positive legs

As *Aedes* spp. of mosquitoes were competent vectors for ILHV following a live feed on a highly viremic (8.18-8.51 log_10_ FFU/ml) mice, artificial bloodmeals were utilized to feed mosquitoes with progressively lower concentrations of virus to determine the oral infectious dose 50% (OID_50_). ILHV FSE 0800 was selected for OID_50_ determination due to its generally higher infectivity in mosquitoes and lower passage history. The OID_50_ in *Ae. aegypti* Salvador was 6.5 log_10_ FFU/ml, and the OID_50_ for *Ae. albopictus* Salvador was 5.5 log_10_ FFU/ml (**Fig. 4A**). To determine whether the susceptibility of the Salvador colonies was broadly representative of *Ae. aegypti* and *Ae. albopictus* or a colony-specific artifact, a geographically diverse panel of *Ae. aegypti* and *Ae. albopictus* were fed artificial bloodmeals with ILHV FSE 0800 at the calculated OID_50_ concentrations (**Fig. 4B-C**). Both species were broadly susceptible across all colonies tested. The eight colonies of *Ae. aegypti* averaged 67% infection (range: 28-100%); the only colony with an infection rate below 50% was Dakar, which was also the lone representative from Africa. The six colonies *of Ae. albopictus* averaged 65% infection (range: 52-74%).

**Fig. 4.**
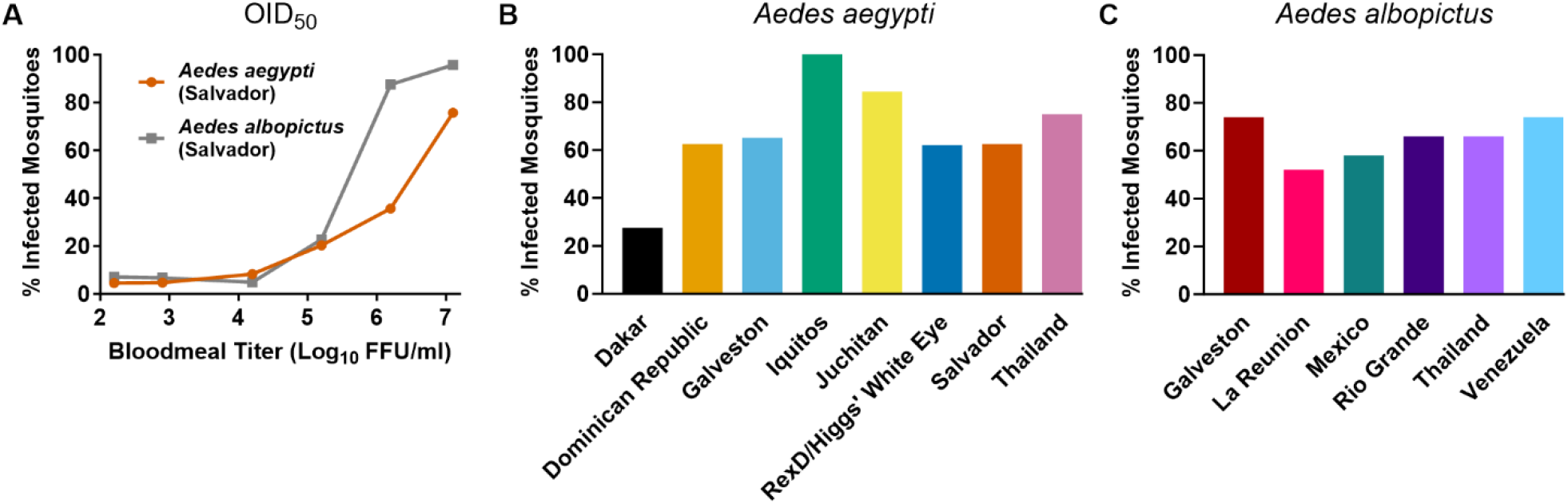
ILHV efficiently infects *Ae*. spp. mosquitoes from a broad geographic range. To determine the OID_50_ of ILHV in *Aedes* spp. mosquitoes, (**A**) *Ae. aegypti* and *Ae. albopictus*, both originating from Salvador, Brazil, were fed artificial bloodmeals containing 2.2-7.1 log_10_ FFU/ml of ILHV FSE 0800. To determine the susceptibility of (**B**) *Ae. aegypti* and (**C**) *Ae. albopictus* from a broad geographic range to ILHV infection, of mosquitoes from various colonies were fed artificial bloodmeals containing ILHV FSE 0800 at the OID_50_ concentration calculated for *Ae. aegypti* (Salvador) and *Ae. albopictus* (Salvador), respectively. In all experiments, engorged mosquitoes were maintained for an EIP of 14 days prior to harvest and the presence or absence of ILHV was determined by infectious assay with immunostaining in Vero cells. OID_50_ values (**A**)were calculated by probit analysis using the rates of infection calculated from an average of 38 (range: 22-59) *Ae. aegypti* and 40 (range: 23-57) *Ae. albopictus* per dose. Rates of infection from geographically diverse *Aedes* spp. mosquitoes (**B-C**) were calculated from an average of 28 (range: 8-40) *Ae. aegypti* and from 50 *Ae. albopictus* per colony. Data are representative of a single experiment.

## Discussion

Although ILHV was discovered in 1944, relatively little is known about this virus outside of serosurveys and case reports from sporadic symptomatic human infections (*1, 8, 11, 13, 20-24, 27, 28, 30, 31, 34, 35, 38, 42-47*). Given that related flaviviruses such as Bussuquara virus, Cacipacoré virus, dengue virus (DENV), Zika virus (ZIKV), West Nile virus (WNV), Saint Louis encephalitis virus (SLEV), yellow fever virus (YFV), and ROCV are endemic throughout the reported geographic range of ILHV (*20, 24, 48, 49*), and the tendency for serological cross reactivity between flaviviruses (*50-53*), the actual disease burden of ILHV is difficult to assess and quantify. Nonetheless, to date, no large-scale human epidemics or epizootic amplifications have been reported. In the past three decades, both WNV and ZIKV have been introduced to the New World and rapidly became endemic (*54-57*). The spread and endemicity of WNV in the Americas underscore the potential for these viruses to rapidly become a serious public health concern out of relative obscurity. Against the landscape of factors such as increasing urbanization of natural biomes, forest degradation for agricultural use, and climate change, the incidence of humans encountering wild animals is ever increasing, thereby increasing the potential for zoonotic spillover events (*40, 58, 59*). Proactively developing a solid knowledge foundation as well as agent-specific reagents and models for under-characterized zoonoses with a strong potential for emergence will allow public health and research efforts more efficiently respond to future outbreaks.

ILHV has been isolated from humans as well as from a variety of bird and mosquito species (*1, 5, 6, 8-17*). However, serologic evidence indicates that a wide range of vertebrates may be susceptible hosts, including additional bird species as well as mammals and reptiles (*7, 16, 18-32*). Such a range would not be unprecedented; WNV has been detected in nearly 300 bird species and over 30 other species including mammals, reptiles, and amphibians (*60, 61*). Combined with the diverse habitats and feeding behaviors of the mosquito species in which the virus has been detected, ILHV may exist in several different transmission cycles. As this knowledge is necessary to efficiently deploy surveillance and preventative measures such as mosquito spraying, we endeavored to characterize ILHV replication in cell lines from a variety of species. We found that all seven cell lines supported robust replication of all eight ILHV strains tested. Increased peak titers in the RNAi-deficient *Ae. albopictus* cell lines (C6/36 and C7/10) compared to the RNAi-intact *Ae. albopictus* cell line (U4.4) suggests that ILHV is indeed sensitive to the RNAi response of mosquitoes; similar RNAi sensitivities have been reported for other flaviviruses including DENV and YFV (*62, 63*). Peak titers were remarkably similar across the vertebrate cell lines despite representing primates, rodents, and marsupials. This suggests that the wide-ranging detection of antibodies against ILHV may be due to productive ILHV infections rather than cross-reactivity following infection with a related flavivirus. Concerted efforts to isolate ILHV or detect ILHV genomes in samples from a variety of species as part of surveillance efforts will be required to confirm this finding and to better define the transmission cycle(s) of ILHV.

Reproducible animal models are necessary to investigate the pathogenesis of emerging agents as well as to evaluate candidate therapeutics and vaccines. We aimed to develop a series models of ILHV infection that (1) recapitulates human disease, (2) does not require adapted ILHV strains, which may preclude studies with more relevant, circulating ILHV strains, (3) does not require transgenic animals with human gene orthologs, which are frequently expensive and may not be readily available to researchers in all regions, (4) does not require genetic or transient immunosuppression, to facilitate vaccine studies and investigations of virus-host interactions which depend upon an intact immune response, and finally (5) produces readily measured phenotypes such as lethality, morbidity, or viremia to evaluate the efficacy of medical countermeasures. We found that CD-1 mice are a desirable model for ILHV infection, and that this model exhibits ILHV strain-specific phenotypes. Infection of young (four-week-old) mice leads to substantial weight loss and uniform lethality following ILHV Original infection, making this older, more highly passaged strain ideal for testing candidate therapeutics and vaccines. Tropism studies revealed high titers of ILHV Original in the brain and spinal cord; this is consistent with the reported neurological symptoms in severe human infections (*11, 13, 38*). The FSE 0800 strain was also neuroinvasive, albeit to a lesser degree than the Original strain, but did not induce significant weight loss or lethality. These divergent strain-based phenotypes may indicate this model’s utility in detecting emergent phenotypes should new mutations or genotypes arise within ILHV. Interestingly, 80% of mice had detectable ILHV FSE 0800 titers in the testes at four DPI, with one measured at 7.6 log_10_ FFU/g. This tropism merits further investigation, particularly considering the sexual transmission and testicular damage caused by the related flavivirus ZIKV (*64-67*). Older (eight-week-old) CD-1 mice succumbed to infection at a rate of 60% following IP inoculation with ILHV Original; this provides a useful model for vaccine efficacy studies, which necessitate the use of older animals to accommodate seroconversion following vaccination. Finally, we characterized ILHV in A129 mice, which are *IFNAR*^-/-^. This model was rapidly and uniformly lethal and consistently developed high levels of viremia. Although immunocompromised, this does provide an extremely stringent model for medical countermeasure testing as well as an important tool for vector competence studies.

It has long been assumed that ILHV is primarily maintained in a transmission cycle between birds and *Culex* spp. mosquitoes. Phylogenetic analyses consistently demonstrate that ILHV is closely related to other flaviviruses such as JEV, SLEV, and WNV that can utilize bird species as amplifying hosts, *Culex* spp. mosquitoes as vectors, and can cause encephalitic disease during spillover infections in humans (*40, 68-71*). Direct evidence supporting a *Culex* spp. driven transmission cycle for ILHV includes the detection of ILHV RNA in *Cx. Melanoconion, Cx. declarator*, and *Cx. portesi* and the isolation of ILHV from *Cx. coronator* and diverse species of birds: the little blue heron (*Egretta caerulea*), the shiny cowbird (*Molothrus bonariensis*), the keel-billed toucan (*Rhamphastos sulfuratus*), and the double-collared seedeater (*Sporophila caerulescens*)(*4, 5, 7, 16, 17*). However, we found that two colonies of *Cx. quinquefasciatus*, an important vector for WNV and SLEV which is found in ILHV-endemic regions (*72*), were largely refractory to disseminated infection by both ILHV Original and ILHV FSE 0800 despite engorging on high titer (7.78-8.49 log_10_ FFU/ml) bloodmeals from viremic mice. The highest rate of infection we detected in the *Cx. quinquefasciatus* Sebring colony was 9%; in contrast, WNV has been reported to infect this same mosquito colony at rates of 91-100% when exposed to comparable titer bloodmeals (*73*). *Cx. tarsalis* was much more susceptible to infection and dissemination, but ILHV was not detected in any saliva samples and *Cx. tarsalis’* geographic range does not overlap with ILHV-endemic areas, raising questions as to its vector potential. Interestingly, *Ae. aegypti* and *Ae. albopictus* were far more susceptible to infection and dissemination to both the legs and saliva. Dissemination was especially pervasive and rapid with the more modern, lower passage FSE 0800 strain of ILHV. Infection and dissemination to the legs by ILHV FSE 0800 was nearly universal by day seven in both colonies of *Ae. aegypti* and *Ae. albopictus*; by day 14, dissemination to the saliva was detected in at least 80% of mosquitoes in all colonies. The high rates of infection in *Aedes*. spp. were widespread amongst colonies from different regions of North America, South America, and Asia; the only colony with modest rates of infection was an *Ae. aegypti* colony from Dakar, Senegal. Additional mosquito colonies from Africa must be tested to determine whether this is a coincidence or an accurate reflection of a New World flavivirus that infects African mosquitoes less efficiently. Vector competence studies are difficult to directly compare across studies due to the potential impacts of the source, generation, rearing of the mosquitoes, the specific composition and source of the bloodmeal, and the viral stock preparation and detection methods. However, our calculated OID_50_ values of 5.5 log_10_ FFU/ml in *Ae. albopictus* and 6.5 log_10_ FFU/ml in *Ae. aegypti* are broadly in line with the approximately 5-6 log_10_ FFU/ml range most commonly reported for DENV and YFV, both of which are demonstrated to be efficiently transmitted to humans by *Aedes*. spp. mosquitoes (*74-78*). Taken in whole, this work raises the possibility that *Aedes*. spp. mosquitoes may play an important role now, or possibly in the future, in ILHV transmission despite previous assumptions that *Culex*. spp. would be the primary ILHV vectors.

This study has several limitations. Cultured cells are imperfect models of viral replication; notably, Vero cells are deficient in the synthesis of interferon (IFN) and C6/36 and C7/10 cells are deficient in RNA interference (RNAi) (*79-82*). However, they remain an important method of evaluating viral replication in a wide variety of species, many of which are not suitable or common laboratory models but may be important to the viral transmission cycle. We utilized cell lines to investigate the range of species that can support ILHV replication; however, our panel was limited and did not include cells derived from neurologic tissue, birds, or non-*Ae. albopictus* mosquito species. While not all species have cell lines available, including additional diverse lines or incorporating primary cells would be desirable in future experiments. Furthermore, replication competence in cell culture does not necessarily indicate replication competence in the species from which those cells were derived. Confirming any species as susceptible to ILHV will require detection of genomic material or, ideally, viral isolation from that species. Our animal model development focused on viral load, morbidity, and mortality; future investigations should directly examine pathology and immune markers. We focused on male mice for model development to examine potential parallels between ILHV and findings of testicular tropism and damage following ZIKV infection; future studies should include female mice to examine any potential sex-based differences in viral replication or pathology. Finally, our vector competence studies focused on *Aedes* spp. and *Culex* spp. mosquitoes, as these are the main vectors for the most well characterized flaviviruses. However, it is possible that ILHV is primarily vectored by a different genus. In particular, ILHV has been isolated multiple times from *Psorophora ferox*, and this species was implicated in the 1975-1976 outbreak of ROCV (*1, 33, 83*). ILHV has also been isolated from *Ae. scapularis* and *Sabethes chloropterus*, important bridge and sylvan vectors, respectively (*8, 9, 84, 85*). Expanding the range of species utilized in vector competence studies would provide a more comprehensive picture of ILHV transmission cycles, and thus shed more light on its potential for emergence.

## Materials and Methods

### Ethics Statement

All animal work was performed in accordance with UTMB policy as approved by the UTMB Institutional Animal Care and Use Committee (IACUC), protocol number 1807054, approved on 08/26/2018.

### Viruses and Cells

Vero (CCL-81, ATCC, Manassas, VA) and BHK (CCL-10, ATCC, Manassas, VA) were grown in Dulbecco’s modified eagle medium (DMEM, Gibco, Grand Island, NY), supplemented with 10% heat-inactivated fetal bovine serum (FBS, R&D Systems, Flowery Branch, A) and 1% penicillin-streptomycin solution (10^4^ U/ml and 10^4^ μg/ml solution, respectively) (PenStrep, Gibco, Grand Island, NY). Huh-7 and C7/10 cells were grown in DMEM supplemented with 10% FBS, 1% PenStrep, 1% MEM non-essential amino acid solution (Sigma, St Louis, MO), and 1% sodium pyruvate solution (Sigma, St Louis, MO). C6/36 cells (CRL-1660, ATCC, Manassas, VA) were grown in DMEM supplemented with 10% FBS, 1% PenStrep, and 10% tryptose phosphate broth solution (TPB, Sigma, St Louis, MO). U4.4 cells were grown in Mitsuhashi and Maramorosch Insect Medium (HiMedia Laboratories, Kennett Square, PA), supplemented with 20% FBS, 1% PenStrep, 5% TPB, and 0.15% sodium bicarbonate (7.5% w/v solution) (Corning, Manassas, VA). OK cells (CRL-1840, ATCC, Manassas, VA) were grown in minimum essential media (MEM, Gibco, Grand Island, NY), supplemented with 10% FBS and 1% PenStrep. Vero, BHK, Huh-7, and OK cells were maintained at 37°C with 5% CO_2_. C6/36, C7/10, and U4.4 cells were maintained at 28°C with 5% CO_2_.

Eight strains of ILHV (**Table S1**) were obtained from the World Reference Center for Emerging Viruses and Arboviruses (WRCEVA) at the University of Texas Medical Branch (Galveston, TX). The viruses were passaged once in Vero cells to generate working stocks. Stocks utilized in OID_50_ experiments underwent one additional passage in C7/10 cells.

### Virus Quantification

ILHV was quantified via the focus forming assay. Briefly, virus was serially diluted in Vero maintenance media (MEM supplemented with 2% FBS and 1% each of GlutaMax (Gibco, Grand Island, NY), PenStrep, and sodium bicarbonate and inoculated onto 12-well plates of Vero cells. Virus was allowed to infect for one hour in a 37°C, 5% CO_2_ incubator. Following this incubation, an overlay of Opti-MEM (Gibco, Grand Island, NY) supplemented with 2% FBS, 1% PenStrep, and 1% carboxymethylcellulose (Sigma, St Louis, MO) was added to the wells and the plates were returned to the 37°C, 5% CO_2_ incubator. After three days, the plates were fixed with 10% buffered formalin. Fixed plates were blocked with 5% non-fat milk in phosphate buffered saline (PBS, Sigma, St Louis, MO). Anti-ILHV mouse immune ascitic fluid (WRCEVA, Galveston, TX) was diluted 1:1000 in blocking buffer and allowed to bind to the fixed monolayers. Plates were washed with PBS, and HRP-conjugated goat anti-mouse IgG secondary antibody (KPL, Gaithersburg, MD) was diluted 1:2000 in blocking buffer and allowed to bind. Plates were subjected to a final set of washes and developed using KPL TrueBlue peroxidase substrate (SeraCare, Milford, MA).

### Replication Kinetics

Comparative multi-step growth curves of all available ILHV strains were performed in triplicate on Vero, BHK, Huh-7, OK, C6/36, C7/10, and U4.4 cells. Cells were seeded one day prior to infection in 12-well plates. Cell lines were infected with ILHV strains at multiplicity of infection of 0.01 in their respective cell culture media and maintained for one hour at the appropriate temperature for the cell line (28°C for mosquito cells, 37°C for vertebrate cells) with 5% CO_2_. Following the one-hour incubation, cells were washed with PBS and fresh media was added with the FBS content reduced to 2%. Aliquots were harvested daily through 11 DPI, at which time they were clarified by low-speed centrifugation and stored at -80°C. Monolayer integrity was documented every other day from days 2-10 post-infection utilizing a CKX53 inverted microscope (Olympus, Waltham, MA) with the 10x CACHN-IPC objective lens (Olympus, Waltham MA) and an attached EP50 digital camera (Olympus, Waltham, MA). Viral titer was determined by focus forming assay.

### Mouse Model Development and Tissue Tropism

To measure survival, male CD-1 mice (Charles River, Raleigh, NC), either four-or eight-weeks-old, were infected with 5.0 log_10_ FFU of either ILHV FSE 0800 or ILHV Original in a 100μl volume either IP or SC in the back. Cohorts of ten (four-week old IP and eight-week-old SC) or five (eight-week-old SC) infected mice and five (four-week old IP and eight-week-old SC) or two (eight-week-old SC) mock-infected mice were observed out to 21 (eight-week-old IP), 24 (eight-week-old SC), or 28 days post-infection (four-week-old IP). Blood was collected retro-orbitally (RO) from half of the mice on days one and three post-infection and from the other half of the mice on days two and four post-infection. Serum was separated from the blood and stored at -80°C. Mice were weighed daily through at least the first 14 days post-infection.

To assess tissue tropism, four-and ten-week-old male CD-1 mice (Charles River, Raleigh, NC) were infected with 5.0 log_10_ FFU of either ILHV FSE 0800 or ILHV Original in a 100μl volume either IP or SC in the back. Cohorts of ten infected mice and four mock-infected mice were all bled RO on day one post-infection. On day two, half of the mice were euthanized for tissue harvest. The remaining mice were RO bled on additional time on day three, then euthanized on day four for tissue harvest. At the time of harvest, blood was collected from a terminal cardiac puncture and the animal was perfused with sterile PBS. Brain, liver, kidney, muscle (rear hamstring), heart, testes, lung, spleen, spinal cord, and eye tissue were harvested in MEM supplemented with 2% FBS and 1% PenStrep and stored at -80°C, along with the serum that was separated from the blood. Tissues were thawed at the time of titration and homogenized for one minute at a frequency of 26 sec^-1^ in a TissueLyser (Qiagen, Haan, Germany) prior centrifugation for 5 minutes at 16,100xg to pellet debris. Focus forming assays were then performed as described above.

### Vector Competence

Mosquito susceptibility to ILHV infection was determined via live feeds on viremic mice. Eleven- to twelve-week-old female A129 mice (bred at UTMB) were infected SC in the back with 5.0 log_10_ FFU of either ILHV FSE 0800 or ILHV Original. At two DPI, the mice were anesthetized with ketamine/xylazine (100mg/kg and 10 mg/kg, respectively) (for *Aedes* spp. mosquitoes) or ketamine/xylazine/acepromazine (100 mg/kg, 10 mg/kg, and 2 mg/kg, respectively) (for *Culex* spp. mosquitoes). A single mouse was used to feed each carton containing either 150 (for *Aedes* spp.) or 100 (for *Culex* spp.) female mosquitoes. The experiment was conducted using *Ae. aegypti* (Salvador and Venezuela), *Ae. albopictus* (Iquitos and Salvador), *Cx. quinquefasciatus* (Salvador and Sebring), and *Cx. tarsalis* (CDC Ft. Collins) mosquitoes (**Table S3**). The mosquitoes were allowed to feed for 15-30 minutes, after which time the engorged mosquitoes were sorted and returned to their cartons to be held at 27°C with 80% humidity and given access to 10% sucrose for the duration of the study. Mice were euthanized immediately following the mosquito feeds, and serum was collected for immediate titration via focus forming assay.

Mosquitoes were collected at both seven- and 14-days post-feed. At the time of harvest, mosquito bodies and legs were collected and placed in homogenizer tubes with a homogenizer bead and 500μl mosquito homogenization media (DMEM containing 2% FBS, 1% PenStrep, and 1% Fungizone (Gibco, Grand Island, NY)), then stored at -80°C until ready for processing. For a subset of five mosquitoes per cohort in the *Aedes* spp. studies and 20 mosquitoes per cohort in the *Culex* spp. studies, saliva was also collected by immobilizing the mosquito with oil on a microscope slide, allowing the mosquito to salivate into a 10μl pipette tip containing 8μl FBS for one hour, and ejecting the saliva/FBS mixture into 250μl mosquito homogenization media to be stored at -80°C until ready for processing.

The presence or absence of ILHV in mosquito bodies, legs, and saliva was determined via a cytopathic effect assay in Vero cells with immunostaining. Mosquito bodies and legs were homogenized for one minute at a frequency of 26 sec^-1^ in a TissueLyser. Samples were centrifuged for 5 minutes at 16,100xg to pellet debris. Finally, the media was removed from 96-well plates of Vero cells and replaced with 50μl of Vero maintenance media supplemented with 1% Fungizone, and 100μl of undiluted sample was added to the well. In addition to the experimental samples, each plate contained virus stocks as positive controls and homogenized, uninfected whole mosquitoes as negative controls. Following infection, the plates were kept in a 37°C, 5% CO_2_ humidified incubator and then fixed and stained as described for the focus forming assay. The presence of any detectable virus led the well in question to be considered positive. A subset of samples was later removed from the freezer for focus forming assays as previously described to quantitate the viral load.

### Mosquito OID_50_ Determination

The OID_50_ of ILHV FSE 0800 was determined in *Ae. aegypti* (Salvador) and *Ae. albopictus* (Salvador) mosquitoes. Artificial bloodmeals were generated by combining serially diluted ILHV FSE 0800 with washed, packed human erythrocytes from whole blood (Gulf Coast Regional Blood Center, Houston, TX) supplemented with FBS, sucrose, and ATP. An aliquot of each bloodmeal was retained for titration simultaneous to feeding. The bloodmeals were presented to cartons of 100 female mosquitoes in warmed Hemotek feeders (Hemotek Ltd, Blackburn, UK) with mouse skins for 30-60 minutes. Engorged mosquitoes were then sorted and maintained as previously described. Fourteen days after feeding on the bloodmeal, whole mosquitoes were collected in a homogenizer tube with 500μl mosquito homogenization media and stored at -80°C until ready for processing as previously described.

Following OID_50_ determination, the susceptibility of a broad range of both *Ae. aegypti* and *Ae. albopictus* mosquito colonies to ILHV FSE 0800 was determined (**Table S3**). Bloodmeals were prepared as before, with the ILHV FSE 0800 concentration set to the previously calculated OID_*50*_ level determined in *Ae. aegypti* (Salvador) and *Ae. albopictus* (Salvador). An aliquot was again retained for titration simultaneous to feeding, which took place as previously described. Fourteen days after feeding on the bloodmeal, mosquitoes were harvested. Samples were again stored at -80°C until ready for processing as previously described above.

### Statistical Analysis

Survival differences for each cohort pair were assessed using the log-rank (Mantel-Cox) test, with the Holm-Sidak correction for multiple comparisons. Weight changes viral titers were analyzed by two-way repeated measures ANOVA with the strain and day as fixed factors and percent weight change or viral titer as the dependent variable. For replication kinetics data, and for murine data when all three cohorts (ILHV Original, ILHV FSE 0800, and mock) had at least one subject, differences between each cohort pair on a given day were assessed with Tukey’s multiple comparison’s test. When only two cohorts had at least one extant subject (4-week-old mice, IP route, days 9-14), cohorts were compared by multiple t-tests with the Holm-Sidak correction for multiple comparisons. OID_50_ values were calculated by probit analysis using SPSS v25 (IBM, Armonk, NY). Values below the limit of detection were treated as one-half of that limit for graphing and statistical purposes. For titers reported as FFU/g, for which the limit of detection is dependent on the exact weight of the collected tissue and is therefore variable from sample to sample, the lowest value for all samples of that tissue type was used as the limit of detection. All titer data was log_10_ transformed prior to analysis to better approximate normality. An alpha of 0.05 was adopted as the cutoff for statistical significance for all tests a priori. Statistical analysis and graphing performed in Prism v10.0.0 (GraphPad, San Diego, CA). Raw data is available in **Table S4**, and the results of statistical analysis are available in **Table S2**.

## Funding

The research was funded in part by grants awarded by the National Institutes of Health, specifically an International Collaborations in Infectious Disease Research (ICIDR) grant U01AI115577 (NV), a Centers for Research in Emerging Infectious Diseases (CREID) “The Coordinating Research on Emerging Arboviral Threats Encompassing the Neotropics (CREATE-NEO)” grant 1U01AI151807 (NV) and the World Reference Center for Emerging Viruses and Arboviruses (WRCEVA) grant R24AI120942 (SCW). MLN is funded by FAPESP (grant # 2022/03645-1). MLN and REM are CNPq Research Fellows.

## Author contributions

Conceptualization: KSP, JAP, SCW, NV

Methodology: KSP, JAP, SRA, NV

Validation: KSP, JAP, NV

Formal Analysis: KSP, JAP, AFV, NV

Investigation: KSP, JAP, SRA, DPS, DS, AFV, NIOdS, TS, LS, EBF, SLR, NV

Resources: KSP, SCW, REM, MLN, NV

Writing—original draft: KSP, JAP, SRA, NV

Writing—review & editing: KSP, JAP, SRA, DPS, DS, AFV, NIOdS, TS, LS, EBF, SLR, SCW, REM, MLN, NV

Visualization: KSP, JAP, SRA, AFV, NV

Supervision: KSP, MLN, NV

Project Administration: KSP, MLN, NV

Funding Acquisition: SCW, MLN, NV

## Competing interests

Authors declare that they have no competing interests. The funders had no role in the design of the study; in the collection, analyses, or interpretation of data; in the writing of the manuscript, or in the decision to publish the results.

## Data and materials availability

All ILHV strains are available from the World Reference Center for Emerging Viruses and Arboviruses (WRCEVA) at the University of Texas Medical Branch under a material transfer agreement (MTA). The raw data is available in Table S4, and results of all statistical analyses are available in Tables S2. Any additional data will be made available upon request.

## Notes

### Competing Interest Statement

The authors have declared no competing interest.

