## Supplemental Figures S1-10 and Tables S1 and S3 for "Potential of Ilhéus virus to emerge"

Kenneth S. Plante *et al.*

**This PDF file includes:**

Figs. S1 to S10
Tables S1 and S3

Tables S2 and S4 are available separately as .xlsx files due to their size

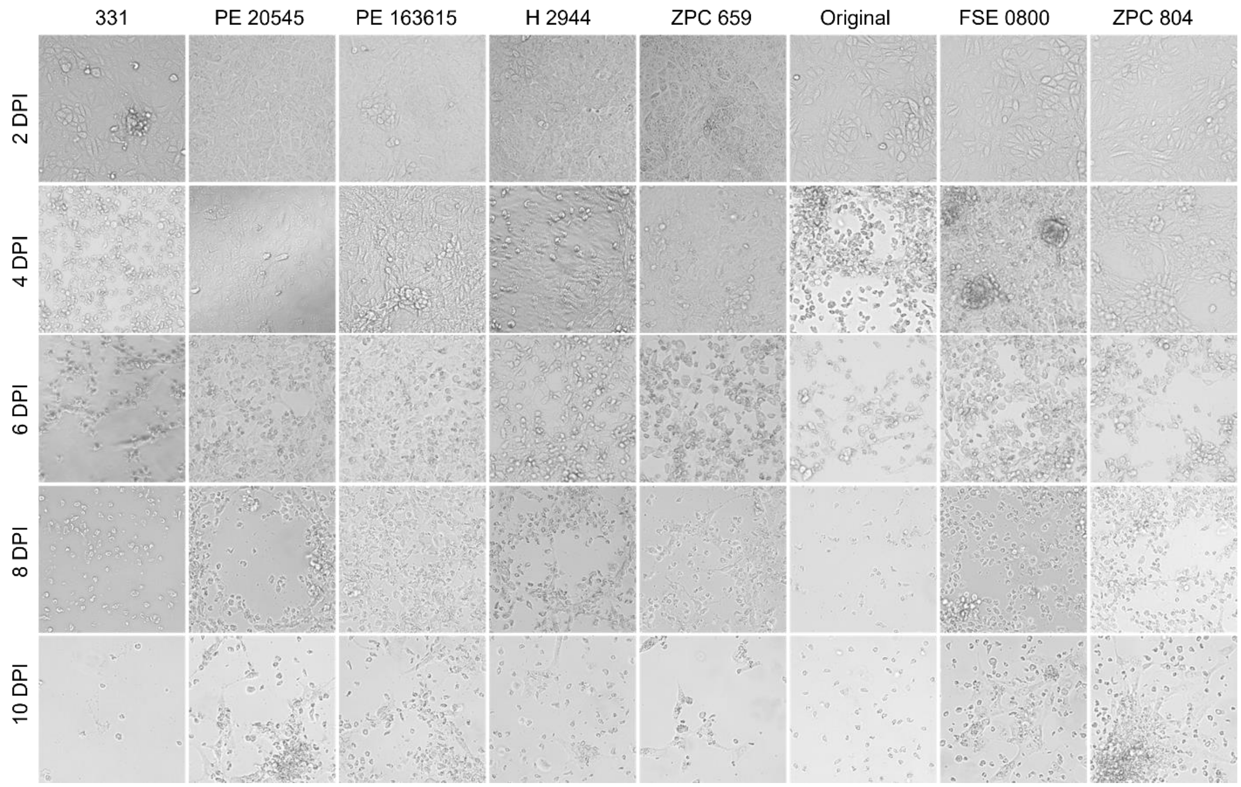


**Fig. S1. Bi-daily monolayer documentation of Vero (*Cercopithecus aethiops*) cells infected with ILHV.** Vero cells were infected with eight strains of ILHV at a multiplicity of infection of 0.01. Monolayer integrity was documented every other day from days two to ten post-infection using a light microscope under 10x magnification.

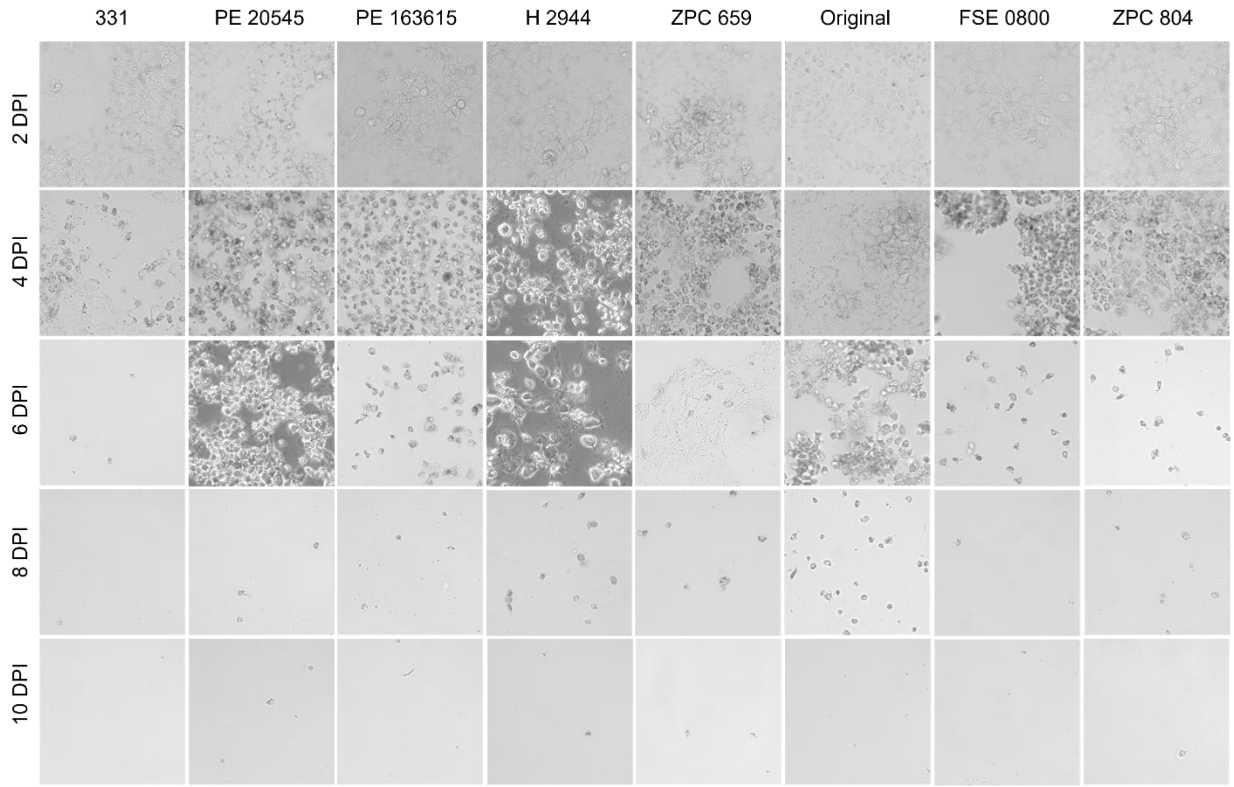

**Fig. S2. Bi-daily monolayer documentation of Huh-7 (*Homo sapiens*) cells infected with ILHV.** Huh-7 cells were infected with eight strains of ILHV at a multiplicity of infection of 0.01. Monolayer integrity was documented every other day from days two to ten post-infection using a light microscope under 10x magnification.

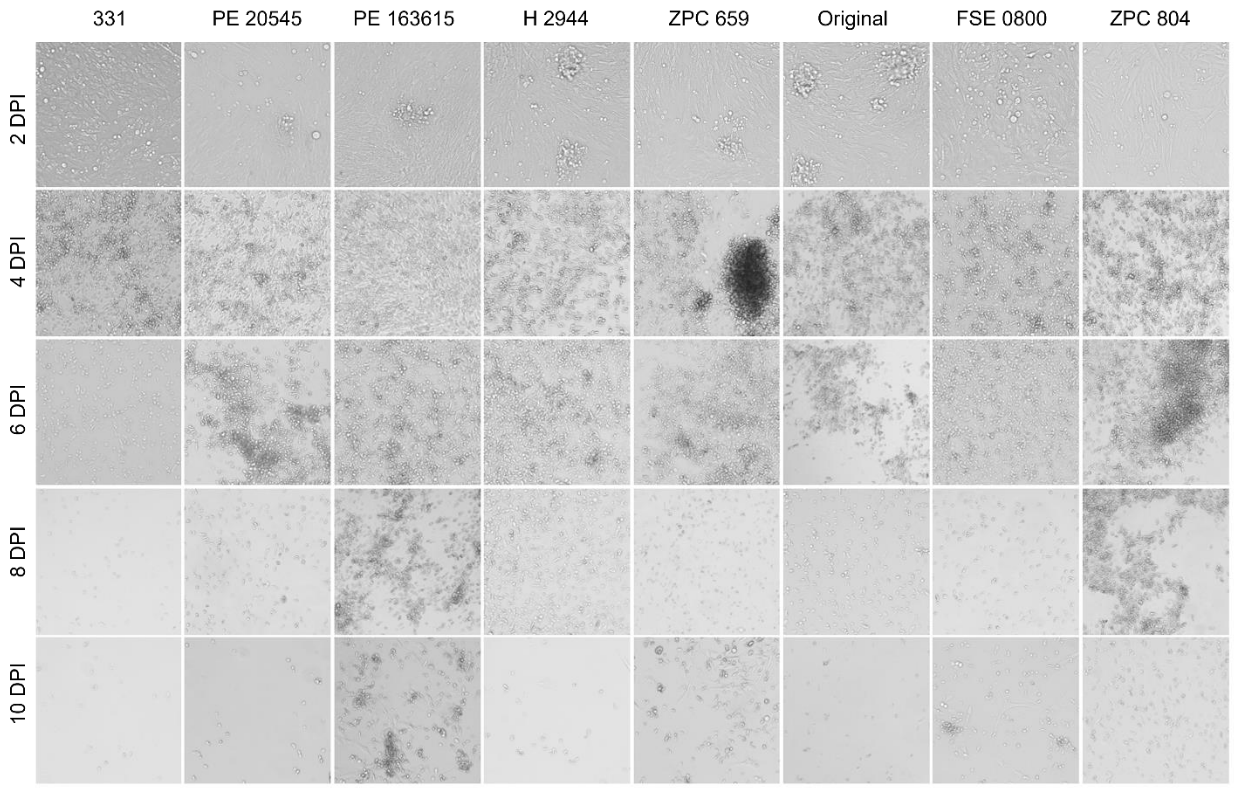


**Fig. S3. Bi-daily monolayer documentation of BHK (*Mesocricetus auratus*) cells infected with ILHV.** BHK cells were infected with eight strains of ILHV at a multiplicity of infection of 0.01. Monolayer integrity was documented every other day from days two to ten post-infection using a light microscope under 10x magnification.

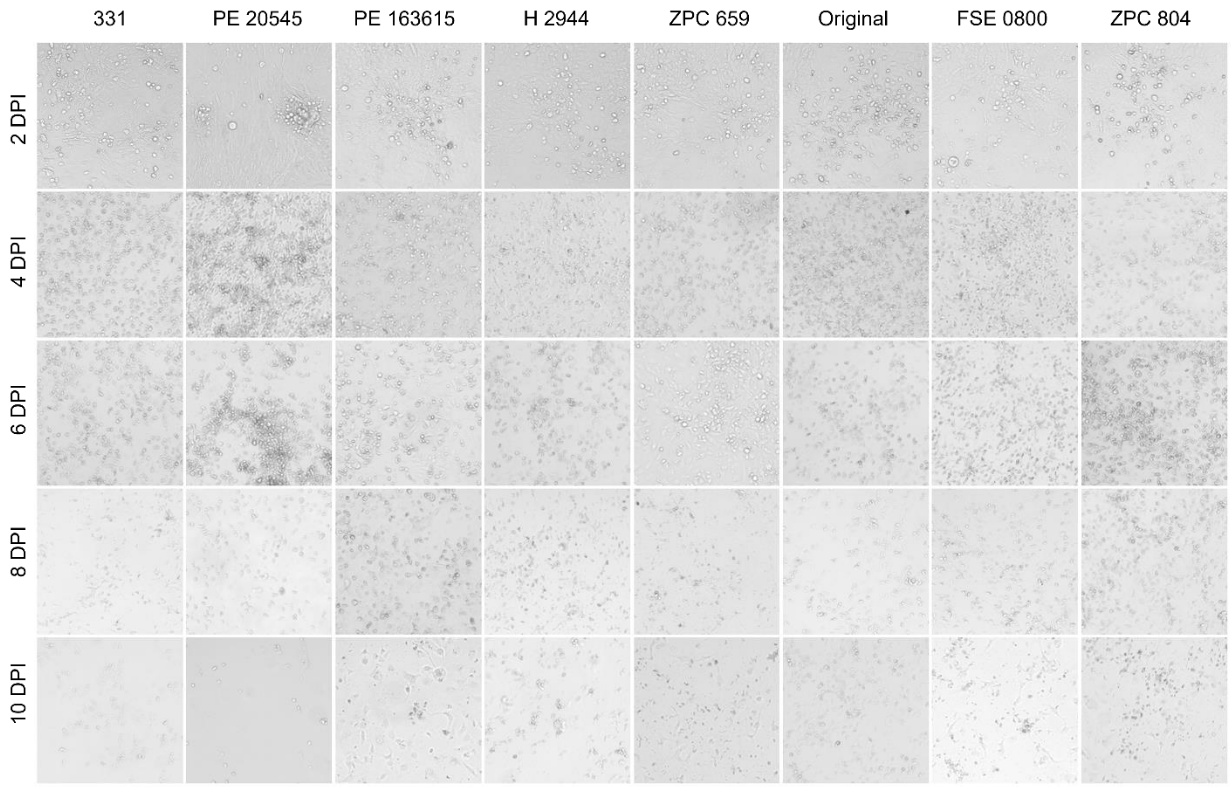


**Fig. S4. Bi-daily monolayer documentation of OK (*Didelphis marsupialis virginiana*) cells infected with ILHV.** OK cells were infected with eight strains of ILHV at a multiplicity of infection of 0.01. Monolayer integrity was documented every other day from days two to ten post-infection using a light microscope under 10x magnification.

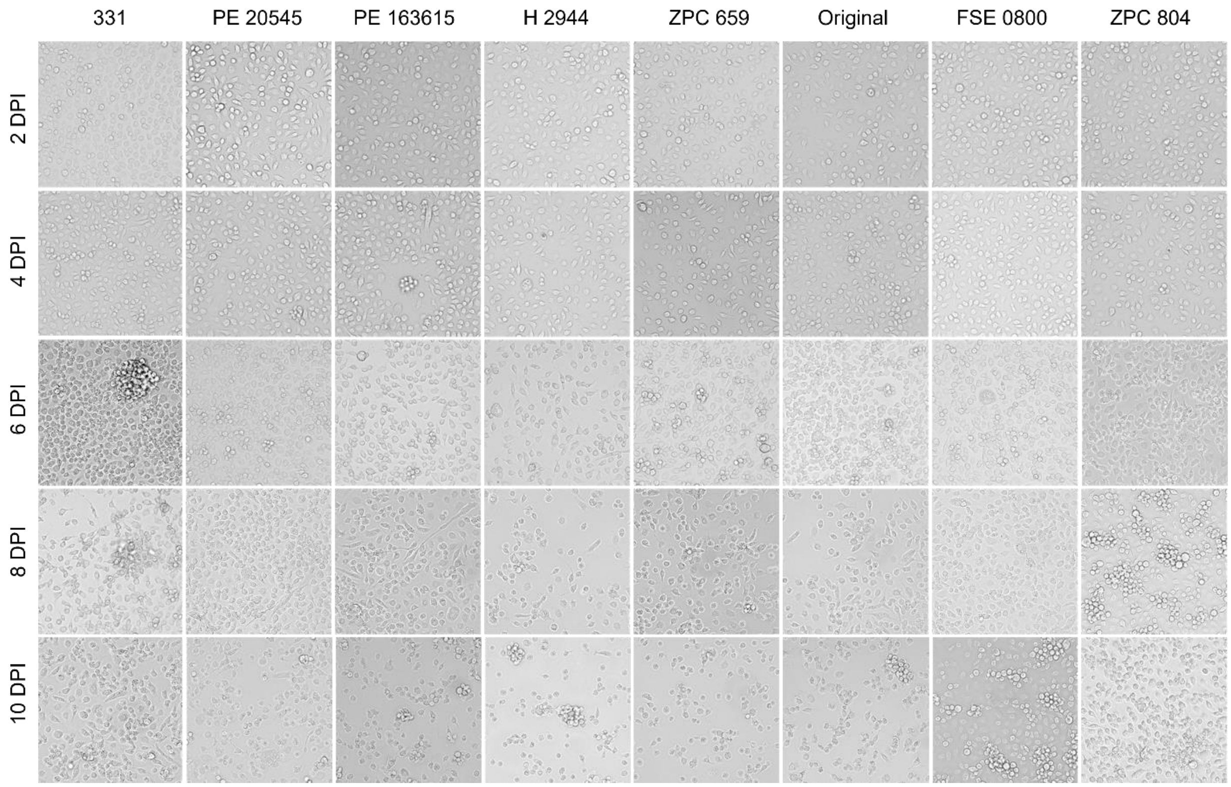


**Fig. S5. Bi-daily monolayer documentation of C6/36 (*Aedes albopictus*) cells infected with ILHV.** C6/36 cells were infected with eight strains of ILHV at a multiplicity of infection of 0.01. Monolayer integrity was documented every other day from days two to ten post-infection using a light microscope under 10x magnification.

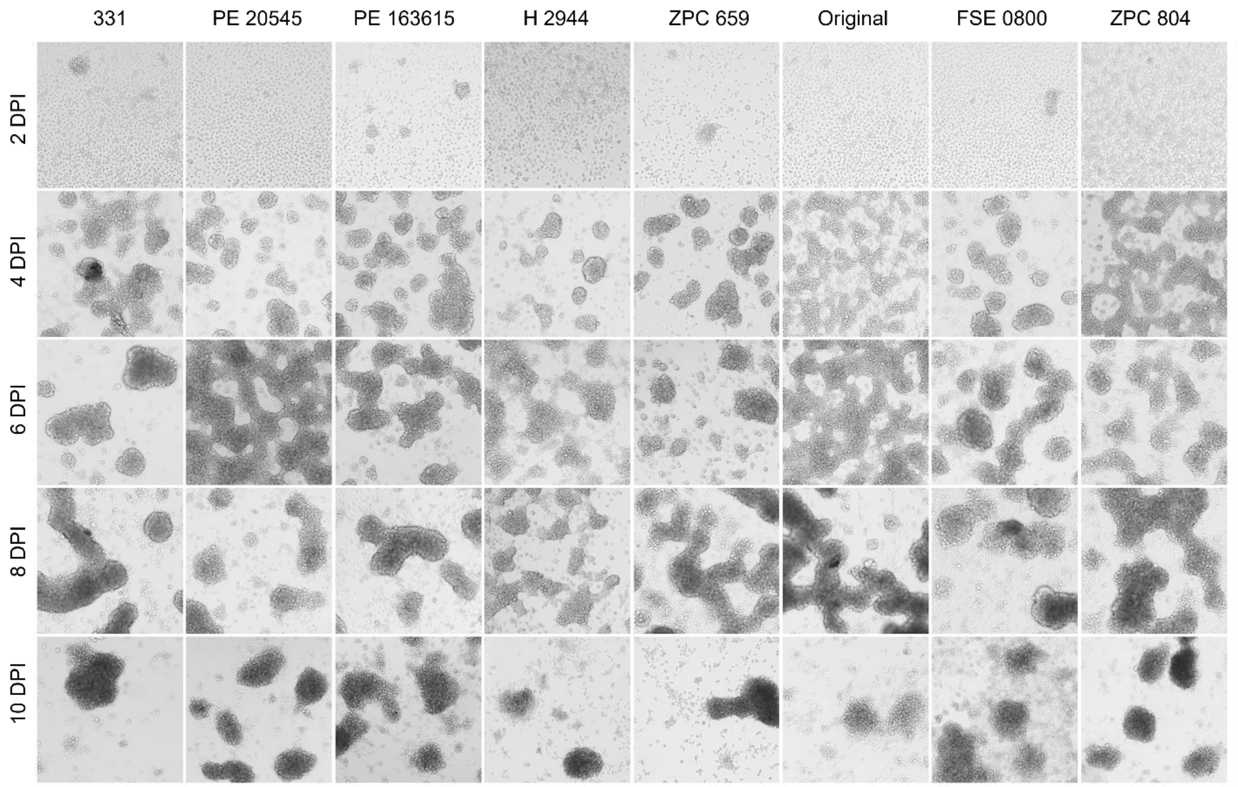


**Fig. S6. Bi-daily monolayer documentation of C7/10 (*Aedes albopictus*) cells infected with ILHV.** C7/10 cells were infected with eight strains of ILHV at a multiplicity of infection of 0.01. Monolayer integrity was documented every other day from days two to ten post-infection using a light microscope under 10x magnification.

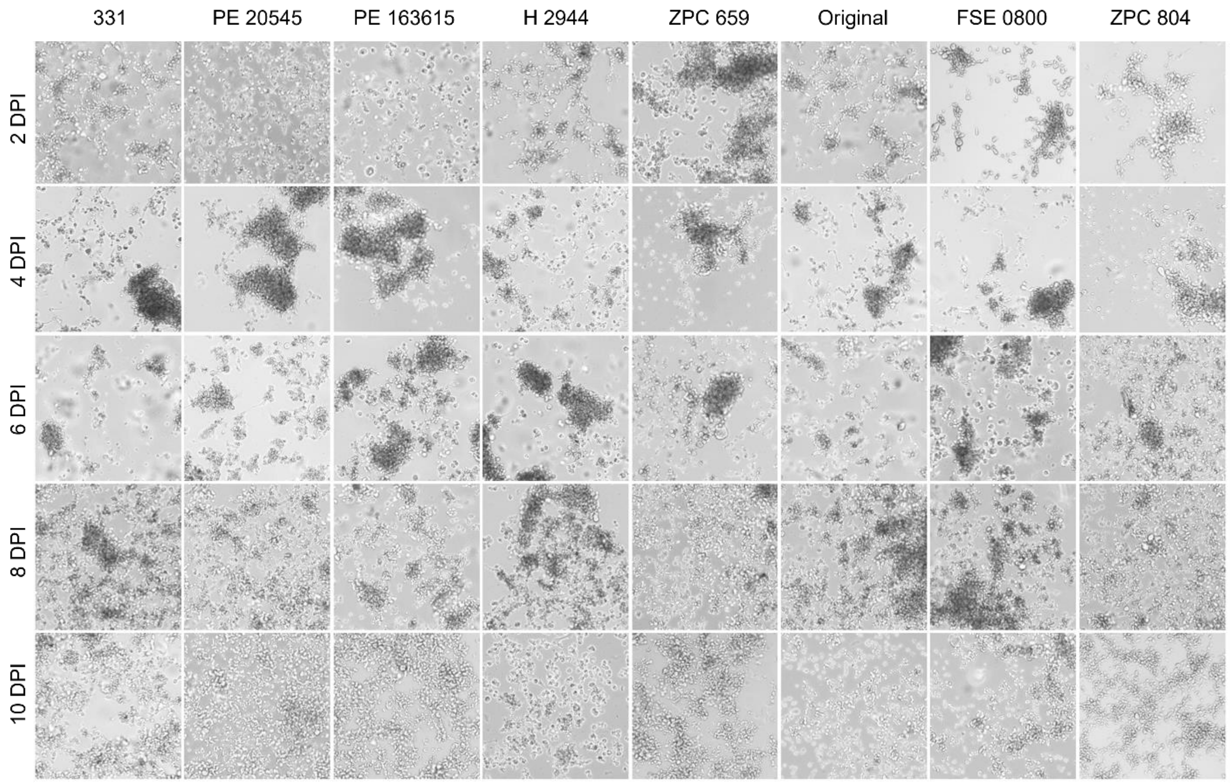


**Fig. S7. Bi-daily monolayer documentation of U4.4 (*Aedes albopictus*) cells infected with ILHV.** U4.4 cells were infected with eight strains of ILHV at a multiplicity of infection of 0.01. Monolayer integrity was documented every other day from days two to ten post-infection using a light microscope under 10x magnification.

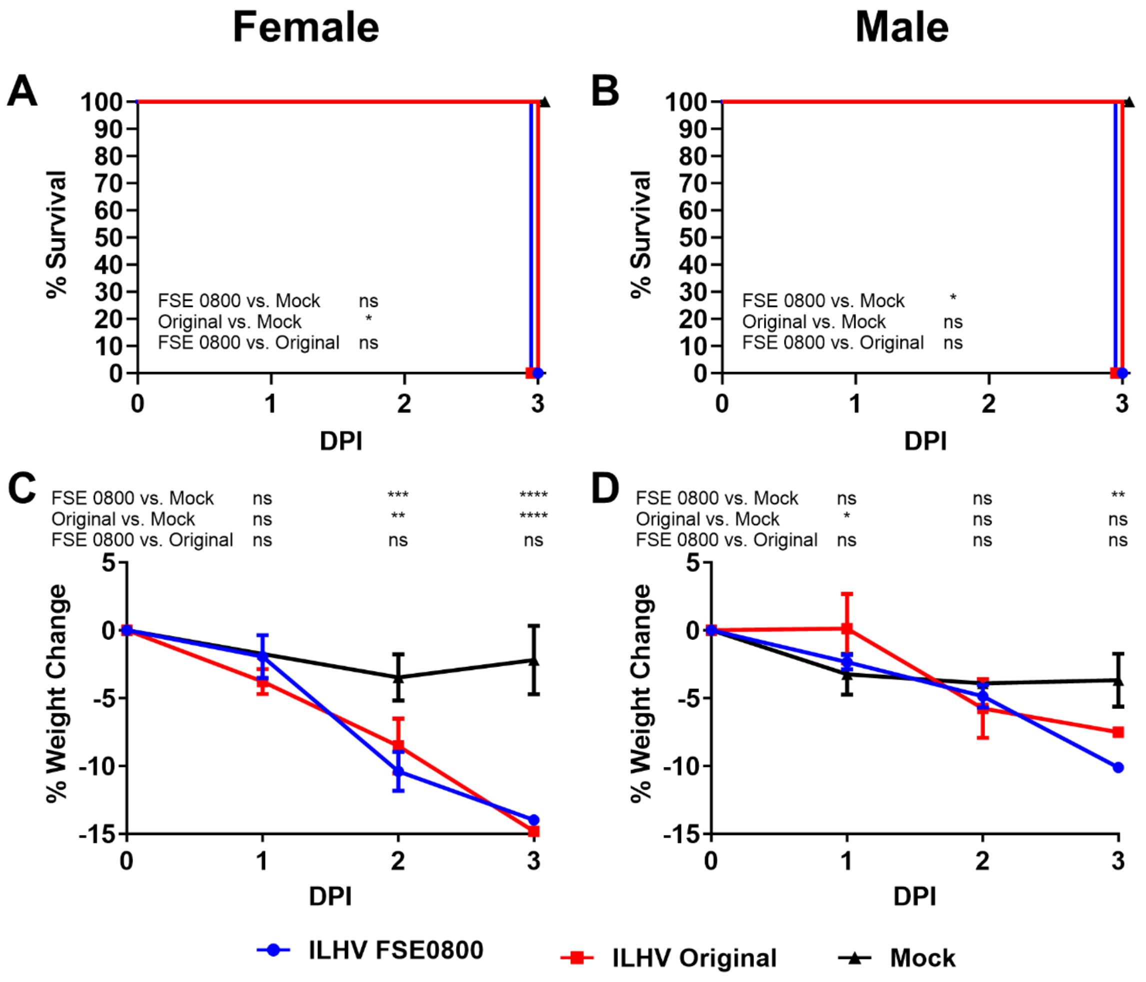


**Fig. S8. ILHV is universally and rapidly lethal in A129 mice.** Six- to eight-week-old, mixed sex, *IFNAR*$^{-/-}$ A129 mice were injected subcutaneously in the back with 5.0 $\log_{10}$ FFU of either ILHV FSE 0800, ILHV Original, or a PBS mock. ILHV FSE 0800 included three females and four males, ILHV Original included four females and three males, and mock included three females and three males. Mice were monitored for survival (**A, B**) and weighed (**C, D**) daily. Survival differences for each cohort pair were assessed using the log-rank (Mantel-Cox) test, with the Holm-Sidak correction for multiple comparisons. Weight changes were assessed by two-way ANOVA with Tukey's test for multiple comparisons. ns = not significant ($p>0.05$), * = $p<0,05$, ** = $p<0.01$, *** = $p<0.001$, **** = $p<0.0001$.

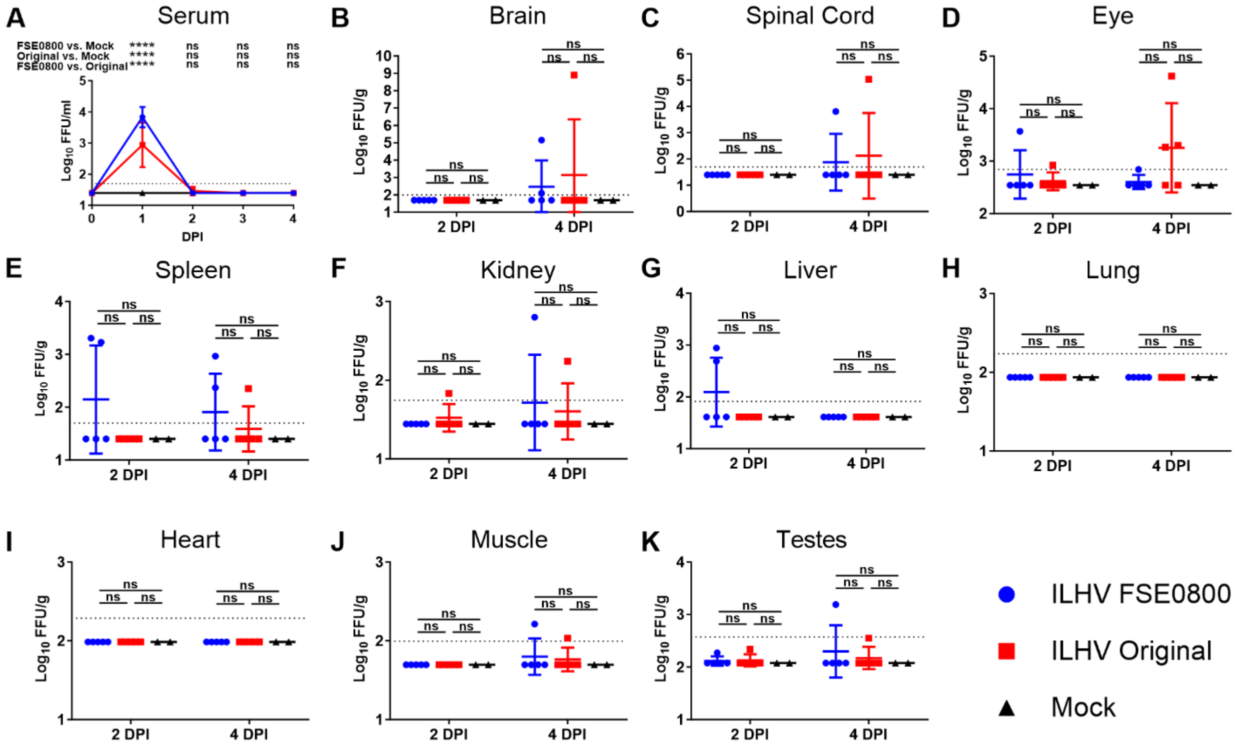


**Fig. S9. Tissue tropism of ILHV in 10-Week-Old CD-1 Mice.** Ten-week-old male CD-1 mice were injected intraperitoneally with 5.0 $log_{10}$ FFU of either ILHV FSE 0800 (n=10), ILHV Original (n=10), or a PBS mock (n=4). Serum was collected retro-orbitally from all mice at one DPI, and from half of the mice at two-four DPI. Half of the mice were euthanized on day two, with the remaining mice euthanized on day four post-infection. Mice were perfused with PBS prior to organ harvest. Titers were determined by focus-forming assay. For the sera (**A**), symbols represent the mean. For the solid organs (**B-K**), symbols represent individual subjects and midlines represent the mean. All error bars represent standard deviation. The dashed black line represents the lower limit of detection. Individuals below the limit of detection are represented as one-half that value for graphing and statistical purposes. Differences between cohort pairs were assessed by Tukey's test for multiple comparisons following two-way ANOVA. ns = not significant (p>0.05), **** = p<0.0001.

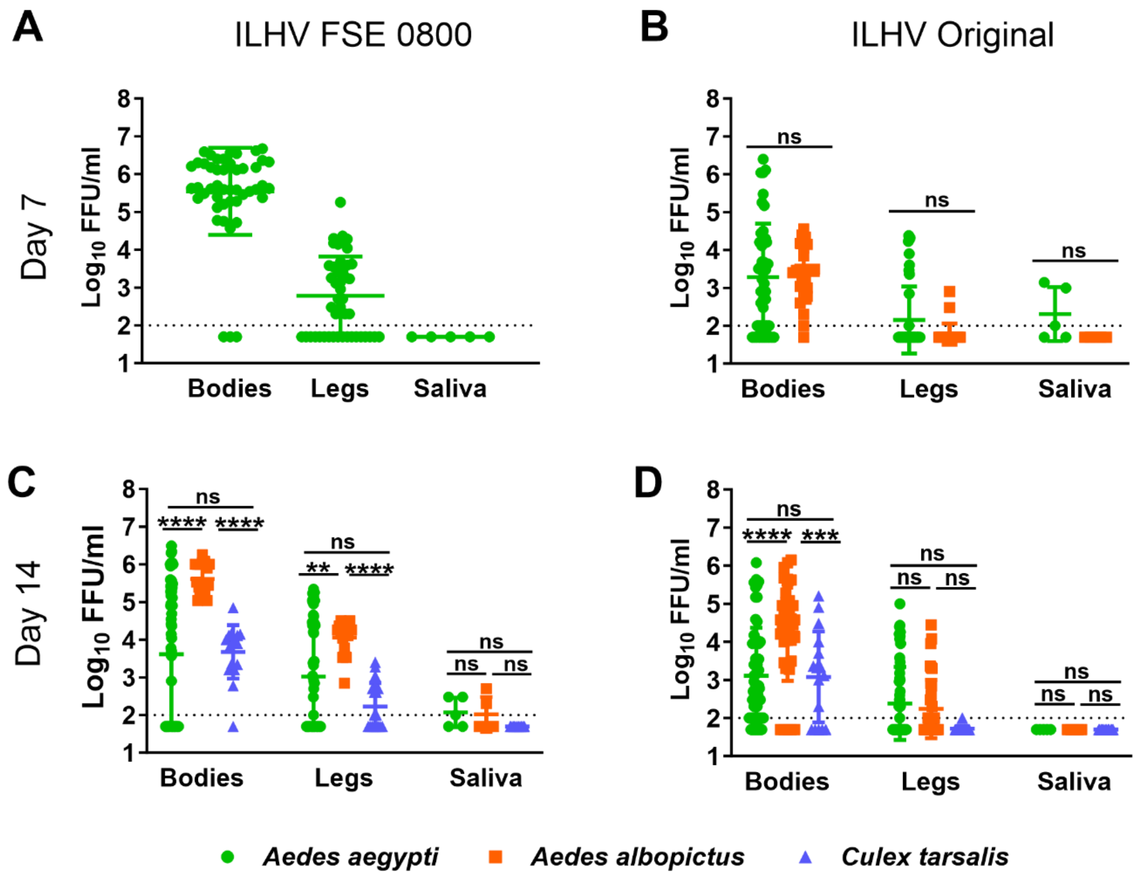


**Fig. S10. Viral Load in the Bodies, Legs, and Saliva of ILHV-Fed Mosquitoes.** *Aedes aegypti* (Salvador), *Aedes albopictus* (Salvador), and *Culex tarsalis* (Ft. Collins) mosquitoes were allowed to feed on viremic A129 mice infected with either ILHV FSE 0800 or ILHV Orignal (mean titer 8.31 $log_{10}$ FFU/ml, range 8.04-8.48 $log_{10}$ FFU/ml). Bodies, legs, and saliva were harvested after an extrinsic incubation period of seven or 14 days. Infectious ILHV titers were determined by focus forming assay, after one previous freeze-thaw to conduct a separate screen for the presence or absence of infectious ILHV. Symbols represent individual mosquitoes, midlines represent the mean, and error bars represent standard deviation. The dashed black line represents the lower limit of detection. Individuals below the limit of detection are represented as one-half that value for graphing and statistical purposes. Statistical analysis was not conducted for ILHV FSE 0800-fed mosquitoes harvested on day seven (**A**) because only one mosquito species was available for analysis. When who mosquito species were available for analysis, as in (**B**), multiple unpaired t-tests were performed with the Holm-Sidak correction. When three mosquito species were available for analysis, as in (**C,D**), differences between pairs were assessed by Tukey's test for multiple comparisons following two-way ANOVA. ns = not significant ($p>0.05$), ** = $p<0.01$, *** = $p<0.001$, **** = $p<0.0001$.

### Table S1. Isolation and Passage History of ILHV Strains

| ILHV Strain | Isolation History | | | | Passage History |
|---|---|---|---|---|---|
| | Source | Country | Location | Date | |
| 331 | Unknown | Brazil | Unknown | 1953 | Suckling Mouse 2, Vero 1 |
| FSE 0800 | Human | Ecuador | Hospital Militar | March 1, 2004 | Vero 4 |
| H 2944* | Mosquito (*Psorophora (Jan.) ferox*) | Peru | Iquitos, Loreto | June 18, 1997 | Vero 5, Hamster 1 |
| Original | Mosquito (*Aedes* and *Psorophora*) | Brazil | Near Ilheus-Fazenda Pirataquiscol | March 6-12, 1944 | Suckling Mouse 29, Vero 2 |
| PE 163615 | Mosquito (*Culex (Mel.) coronator*) | Peru | Iquitos, Loreto | 1999 | Vero 4 |
| PE 20545 | Mosquito (*Psorophora (Jan.) ferox*) | Peru | Iquitos, Loreto | June 18, 1997 | Vero 4 |
| ZPC 659 | Hamster (*Mesocricetus auratus*) | Venezuela | Zulia | 1997 | Vero 2 |
| ZPC 804 | Hamster (*Mesocricetus auratus*) (heart) | Venezuela | Las Nubes, Jesus Ma. Sempum (Catatumbo), Zulia | December 15, 1997 | C6/36 2, Vero 1 |

Passage histories reflect the viral stocks utilized all studies other than $OID_{50}$ experiments, with the final passage being performed in Vero cells. For $OID_{50}$ experiments, one additional passage was performed in C7/10 cells.

*Strain H 2944 is a higher passage derivative of strain PE 20545

**Table S3. Mosquito Colonies.**

| Name | Genus | Species | Location | Generation |
|---|---|---|---|---|
| Dakar | *Aedes* | *aegypti* | Dakar, Senegal | Unknown |
| Dominican Republic | *Aedes* | *aegypti* | Dominican Republic | Unknown |
| Galveston | *Aedes* | *aegypti* | Galveston, TX | Unknown |
| Iquitos | *Aedes* | *aegypti* | Iquitos, Peru | Unknown |
| Juchitan | *Aedes* | *aegypti* | Juchitan, Oaxaca, Mexico | Unknown |
| RexD/Higgs' white eye | *Aedes* | *aegypti* | Rexville, Puerto Rico | Unknown |
| Salvador | *Aedes* | *aegypti* | Salvador, Brazil | F10 |
| Thailand | *Aedes* | *aegypti* | Bangkok, Thailand | Unknown |
| Galveston | *Aedes* | *albopictus* | Galveston, TX | Unknown |
| La Reunion | *Aedes* | *albopictus* | La Reunion | Unknown |
| Mexico | *Aedes* | *albopictus* | Mexico | Unknown |
| Rio Grande | *Aedes* | *albopictus* | Rio Grande valley, TX | Unknown |
| Salvador | *Aedes* | *albopictus* | Salvador, Brazil | F10 |
| Thailand | *Aedes* | *albopictus* | Bangkok, Thailand | Unknown |
| Venezuela | *Aedes* | *albopictus* | Venezuela | Unknown |
| Salvador | *Culex* | *quinquefasciatus* | Salvador, Brazil | Unknown |
| Sebring | *Culex* | *quinquefasciatus* | Sebring, FL | Unknown |
| CDC Ft. Collins | *Culex* | *tarsalis* | Unknown | Unknown |
